# Unveiling Cerebrospinal Fluid Protein Biomarkers in Pediatric Acute Lymphoblastic Leukemia Using Proximity Extension Assay

**DOI:** 10.64898/2026.07.03.736065

**Authors:** Mahshid Moballegh Nasery, Réka Gergely, Nóra Kutszegi, Istvan Szegedi, Dániel János Erdélyi, Csongor Kiss, Éva Csősz

## Abstract

**Background:** Acute Lymphoblastic Leukemia (ALL) is a highly heterogeneous pediatric malignancy. Despite high survival rates, relapse and the involvement of central nervous system (CNS) remains a significant clinical challenge. Traditional clinical parameters often lack the precision required for early detection and risk stratification. This study utilizes high-throughput proteomics and machine learning (ML) to identify molecular signatures in cerebrospinal fluid (CSF) that characterize disease effect and treatment response.

**Methods:** 82 CSF samples from 41 pediatric ALL patients at diagnosis (VD) and remission (VR) were analyzed. Proteomic profiling of 276 proteins was performed using Olink’s Proximity Extension Assay. Differentially abundant proteins were identified (q-value< 0.05, |Log_2FC| > 0.5) using the Wilcoxon rank-sum test. Three ML algorithms - Random Forest, LASSO, and SVM-RFE - were integrated to select the differentially abundant proteins in VR and VD and between CNS involvement levels. To validate the data Pan-Cancer Atlas analysis was done using two different platforms.

**Results:** In VR, we observed significant alterations in the expression of key proteins compared to VD, with ADGRG1 and KYNU showing a marked increase, while CCL17, CD5, CD27, CXCL9, CXCL11, FASLG, GZMA, and TNFRSF9 were significantly downregulated. Furthermore, our analysis identified distinct protein signatures associated with CNS involvement: CCL4, CTSC, CXCL10, CXCL9, and MMP7 were differentially abundant at the VD, whereas CAIX, CASP-8, HAGH, CXCL9, MMP7, MCP-2, and VWC2 at the VR.

**Conclusion:** Integrating Olink proteomics with ML identified molecular signatures in ALL that have the potential to be further developed to a biomarker panel for monitoring treatment response and guiding personalized therapeutic strategies shifting the focus toward the Precision One Health approaches.

## Introduction

Acute lymphoblastic leukemia (ALL), which originates from B- or T-cell lymphoid precursors, is the most common cancer of childhood and adolescence [1]. The disease is not a single entity, but a heterogeneous collection of more than 30 distinct genetic subtypes [2].

Due to remarkable advances in contemporary treatment protocols, five-year survival rates in high-income countries have reached over 90% [3]. However, despite improved survival rates, relapse still occurs in 10 to 15 percent of patients, and death from relapsed ALL remains a leading cause of cancer-related mortality in children [4].

Central nervous system (CNS) involvement is a significant challenge in ALL. While CNS involvement is rare at the time of initial diagnosis (3–5%), it is much more common at relapse, occurring in approximately 30–40% of cases [5]. Although therapeutic advances have minimized relapse outside the nervous system (such as the testes), CNS relapse still occurs in 3–8% of children with ALL. Therefore, there is a need for novel approaches that go beyond classical clinical and laboratory parameters to investigate biological mechanisms in more depth [4].

Proteomics has emerged as a new frontier for understanding the biology of the human body. Since proteins are the ultimate executors of genetic instructions and are responsible for cellular processes, their study bridges the gap between the genome and phenotype. The discovery of protein biomarkers plays a role in early diagnosis and a more accurate prediction of the risk of recurrence [6], [7].

However, proteome analysis in complex biological samples has been accompanied by technical challenges. The emergence of advanced technologies such as proximity expansion assay by the Olink platform, combining the features of immunoassay and the high sensitivity of DNA technologies, has taken a major step in high-throughput and accurate analysis of proteins with a wide concentration range, allowing for accurate molecular mapping [8], [9], [10]. Ultimately, the sheer volume and complexity of data generated by omics technologies are beyond the power of human analysis and intuition. This is where ML come in as strategic tools. Using computational models and ML algorithms, hidden patterns in protein interactions can be identified [9].

This study leverages Olink’s proteomics data and the ML analysis to identify molecular signatures associated with disease progression and treatment response in pediatric patients with ALL.

## Material and methods

### Sample collection

In this study, 82 cerebrospinal fluid (CSF) samples from 41 patients with ALL were taken at diagnosis (VD) and in remission (VR), where the blast cell ratio in bone marrow was <1% [11], using a lumbar puncture procedure. Samples were collected from children with B-cell or T-cell acute ALL who were admitted to the Department of Pediatrics University of Debrecen, and Pediatric Center Semmelweis UniversityThe diagnosis of the disease was confirmed by an oncologist for all patients based on standard criteria, defined as having greater than 20% blast cells in the bone marrow sample at VD [11]. The median age of the patients was 5 years (range 3 to 18 years) and the female: male ratio was 1.15:1 **(Supplementary table 1)**.

CSF samples were collected in sterile tubes and processed within 30 minutes. To separate cells and cellular debris, samples were centrifuged at room temperature at 2500 rpm for 10 min. The supernatant was then removed and aliquot. All samples were stored at −80 °C until analysis.

CNS classification was performed by cytomorphology and/or clinical findings (imaging and biopsy): (1) CNS1, absence of blasts on cytospin preparation in CSF and no clinical or imaging findings of CNS leukemia; (2) CNS2, #5/mL nucleated cells in CSF, cytospin preparation positive for blasts, and no clinical or imaging findings of CNS disease; (3) CNS3: .5/mL nucleated cells in CSF and cytospin preparation positive for blasts or clinical or imaging findings of CNS disease. Presence of leukemic lymphoblasts in the CSF samples was investigated by cytomorphology and confirmed by immunophenotyping with flow cytometry, which helped in defining the presence of leukemic lymphoblasts in the CSF sample in cases that were dubious by morphological evaluation [12], [13], [14].

This study was approved by the Ethics Committee of University of Debrecen, Ethics Code:DEOEC RKEB/IKEB: 6126-2022 on August 02, 2022, and by the national Scientific Research end Ethics Committee (ETT-TUKEB), Ethics code: BMEÜ/1916-4/2022/EKU and approved by the National Centre for Public Health and Pharmacy on February 09, 2023, and the modification was approved by ETT-TUKEB, Ethics code: BM/2373-1/2023 on September 06, 2023, and approved for Semmelweis University with Ethics Code: BMEÜ/3798-1/2022/EKU on November 30, 2022. All participants agreed in participation and the guardians of the donors gave written informed consent. This research was conducted ethically in accordance with the World Medical Association Declaration of Helsinki.

### Proteomics Analysis

The Immuno-oncology, Neurology, and Organ damage Olink Target 96 panels (ThermoScientific) were used to measure the relative amount of 92 proteins per panel in each of the 82 samples. The sample preparation and measurements were performed in our laboratory, and the protocols provided by Olink were rigorously followed [8]. The determination of the normalized protein expression (NPX) values were done by Olink NPX Signature 2.0.2 software following the data acquisition on Olink Signature 100 (ThermoScientific) instrument.

In the quality control phase, one sample did not pass the strict quality control of Olink panels and was excluded from the study. Those proteins, which were either not detected or the NPX value was below limit of detection in more than 70% of the samples, were removed. For the remaining proteins, the initial NPX value was kept, or if the value was below the LOD, the LOD value vas taken. After filtering, 167 proteins were selected for further analysis. No further filtering or imputation was performed.

### Statistical Analysis

Statistical analyses and data visualization were performed using R version 4.5.1. The log2 fold change (FC) was calculated in each case, and the nonparametric Wilcoxon rank-sum test was used to identify differentially abundant proteins. To control the false discovery rate resulting from repeated testing on 167 proteins, p-values were adjusted using qvalue package (Storey, 2002), and differentially abundant proteins were identified based on a q-value. The threshold for statistical significance in all analyses was q < 0.05 and |Log2FC| > 0.5. The difference in protein amounts between the visits (VR vs. VD), type of disease (T-ALL vs. B-ALL), gender (female vs. male), as well as CNS involvement (groups 1, 2, and 3) were examined. For visualization of the results, volcano plots, heatmaps, and boxplots were used. The graphs were drawn using the ggplot2, ggrepel, and pheatmap packages.

### Machine learning

The application of Machine learning (ML) algorithms was performed based on in-house developed scripts with minor changes. Briefly, to identify the proteins that are able to discriminate between the groups, three different feature selection algorithms were used, including Random Forest (RF), Least Absolute Shrinkage and Selection Operator (LASSO), and Support Vector Machine Recursive Feature Elimination (SVM-RFE). In the RF algorithm, proteins were ranked based on the Mean Decrease Gini score to identify the top features and to identify the complex and nonlinear relationships between proteins. Besides RF, LASSO regression with L1-regularization penalty was used. In the SVM-RFE algorithm, a radial basis function was used instead of the linear kernel used in the original code to better identify nonlinear relationships between proteins, and was tuned using the grid search, with parameters included: cost (C): 0.1, 1, 10, 100, and gamma: 0.001, 0.01, 0.1, 1. For each algorithm, a cross-validation (CV) process was implemented to assess the stability of the models. For the RF model, a 5-fold CV structure was designed manually, for LASSO, the built-in capabilities of the glmnet (with 10 iterations), and for SVM-RFE, the caret (with 5 iterations) packages were used to automatically perform CV and parameter optimization. For comparisons regarding the CNS involvement levels, the Leave-One-Out CV was applied.

### Protein Feature Selection

To further examine the proteins selected by the ML models, a two-step procedure was applied. The SVM-RFE algorithm was used, and first, the performance of the proteins selected by all ML models was examined. Then, we added to the model one by one each protein identified by at least one ML algorithm, and those that improved model performance were kept. Finally, different combinations of well-performing proteins were tested. The model performance was examined on Out-of-Bag data, and the different protein combinations were ranked based on the area under the curve (AUC). Those proteins were selected that provided the highest AUC.

### Network and functional analysis

Proteins identified through statistical and ML evaluation were subjected to network and functional analyses. The set of proteins having the highest discriminatory performance was subjected to network analysis. The STRING database (version 12.0) [15] was used, and the protein-protein interactions of the selected proteins and their first shell interactors were visualized, considering the highest confidence score of 0.9. To better understand the biological role of these proteins, functional enrichment analysis was performed.

### External data evaluation

To evaluate the expression profiles of the identified candidates, we utilized the R2: Genomics Analysis and Visualization Platform (http://r2platform.com/TPCPA) [16]. The Pan-Cancer Proteome Atlas was used to assess the expression of these 10 proteins on a larger scale. Analyses were performed in the Gene vs Track format, and one-way ANOVA was used to statistically compare protein expression across different cancer categories. A p-value of less than 0.05 was considered significant, and the data were visualized in the form of box plots.

Survival analysis was performed using the PedcBioPortal for Childhood Cancer Genomics (https://pedcbioportal.org/) [17], [18], [19], utilizing data from the Open Pediatric Cancer (OpenPedCan) Project (v15) [20], To gain a comprehensive understanding of how these gene expression profiles impact patient outcomes, we evaluated their prognostic value across two distinct settings. First, we analyzed the entire, unfiltered pan-cancer dataset (15856 patients) to capture broad clinical trends. Next, we narrowed our focus exclusively to the 1101 patients of the ALL cohort, which provided a targeted sub-cohort of 1,483 samples. For all analyses, we focused on RNA expression z-scores to assess baseline expression levels. Guided by our experimental laboratory findings, we executed two separate query sessions for each cohort: one tracking ADGRG1 and KYNU, and another for the eight genes (CCL17, CD5, CD27, CXCL9, CXCL11, FASLG, GZMA, and TNFRSF9) based on their abundance changes in our dataset. The platform’s automated stratification divided samples into altered and unaltered groups depending on the presence of genomic alterations in the queried genes. We then generated Kaplan-Meier survival curves to estimate Overall Survival. To determine whether the survival differences between these groups were statistically meaningful, a log-rank (Mantel-Cox) test provided by the platform, using a standard significance threshold of p < 0.05, was used.

## Results

High-throughput proteomic profiling of 82 samples was performed using the Immuno-oncology, Neurology, and Organ damage Olink Target 96 panels, tracking 92 proteins per panel. Following data acquisition on the Olink Signature 100 platform and initial processing via Olink NPX Signature 2.0.2 software, one sample failed the quality control criteria and was excluded, leaving 81 samples for analysis. In case of proteins where the NPX was lower than the LOD value, the LOD values were considered and those proteins where the LOD value was present in more than 70% of the samples were removed. No additional filtering or imputation was performed. Ultimately, 167 proteins passed QC and filtering and were selected for subsequent analysis **(Supplementary table 2**).

To investigate overall changes in protein abundance in CSF between diagnosis and remission, NPX values were compared between VD and VR in patients with ALL.

After performing the nonparametric Wilcoxon test, proteins were selected based on a significance threshold of q<0.05 and an effect size of Log2FC>0.5.

In total, of the 167 proteins examined, three proteins (CXCL9, CD27, and GZMA) showed a statistically significant decrease in abundance in remission **(Figure 1A)**, and the pattern of individual responses showed gender-related heterogeneity **(Figure 1B)**.

**Figure 1.**
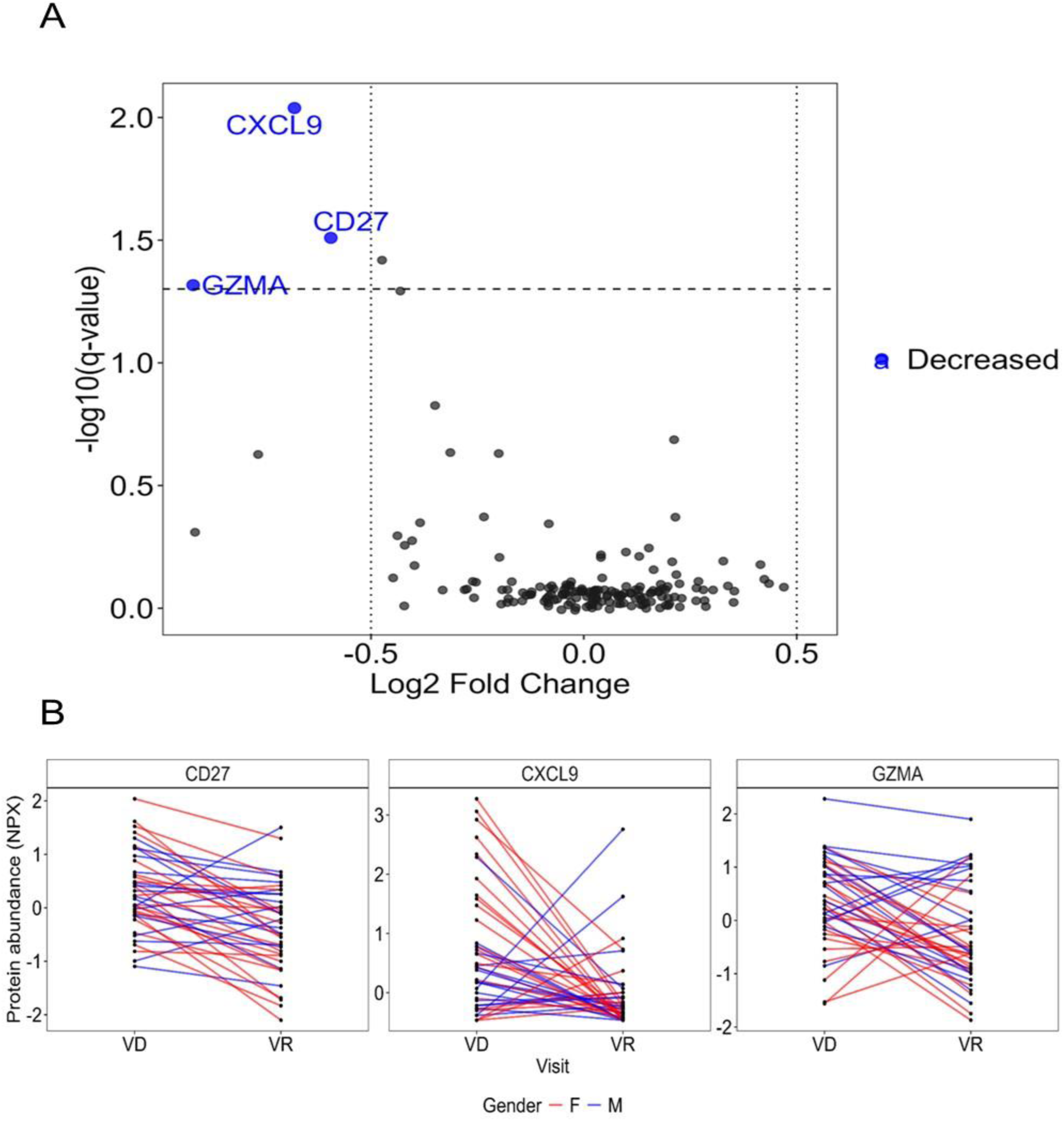
Comparative analysis of protein abundance in ALL patients at VD and VR. (A) Volcano plot shows the distribution of 167 proteins examined based on q-value and Log2FC. The proteins, which are decreasing in VR, are shown in blue. (B) Trajectory plot showing the overall trend of protein abundance change in each case. Red and blue lines indicate female and male patients, respectively. F: female, M: male, VD: visit at diagnosis, VR: visit at remission

The trajectory plots for CD27 and CXCL9 point toward a gender bias. To examine the possible effect of gender on the abundance patterns of the proteins, subgroups were created based on gender, and the statistical analysis was carried out. The results showed that in females but not in males, the difference in the case of these two proteins was statistically significant between the VD and VR **(Supplementary Figure 1)**.

In the case of some patients, a CNS involvement can be observed, and according to the level of infiltration, the patients were grouped into three categories [11]. The CNS1 group was characterized by the absence of blasts in CSF on cytospin preparations and no clinical or imaging findings of CNS leukemia; in the CNS2 group, the cytospin preparations were positive for the presence of blasts (<5/mL nucleated cells) but no clinical or imaging findings of CNS disease was present; while in case of CNS3 group, the cytospin preparations were positive for the presence of blasts (>5/mL nucleated cells) and the clinical or imaging findings were positive for CNS leukemia [13].

In case of one patient, the information on the CNS involvement level was missing, so the two samples of this patient were removed from the analysis. The analysis of patients belonging to each CNS group led to the identification of four proteins in the CNS1 group as differentially abundant between the VD and VR groups, but no statistically significant difference in the CNS2 or CNS3 groups could be observed. All four proteins had a decrease in abundance in the VR group compared to VD **(Supplementary Figure 2)**.The comparison of CNS1 to CNS2 and CNS3, as well as of CNS1 to CNS2+3 both in VD and VR did not show any statistically significant change **(Supplementary Figure 3)**.

In addition, there was no significant difference between VD and VR in the case of B-ALL or T-ALL. Comparison of the disease types revealed that CXCL11, CD5, CD8A, VEGFR2, and ADA were differentially abundant between B-ALL and T-ALL during the VD stage. No Differentially abundant proteins could be identified between the other groups **(Supplementary Figure 4)**.

### Application of Machine Learning algorithms for the identification of group-discriminatory proteins

Three different ML models, RF, LASSO, and SVM, were used to examine the most important proteins able to discriminate among the groups. These multiple approaches were adopted to reduce bias and ensure the selection of features with high accuracy.

In the RF model, the importance of the features was evaluated based on the Mean Decrease Gini score. The results showed the proteins that play an important role in differentiating the VD and VR groups are CXCL9, GZMA, KYNU, FASLG, Gal_9, CD27, CCL3, CCL17, CD5, and CXCL11. The model was able to discriminate properly between the two groups **(Figure 2A)**.

**Figure 2.**
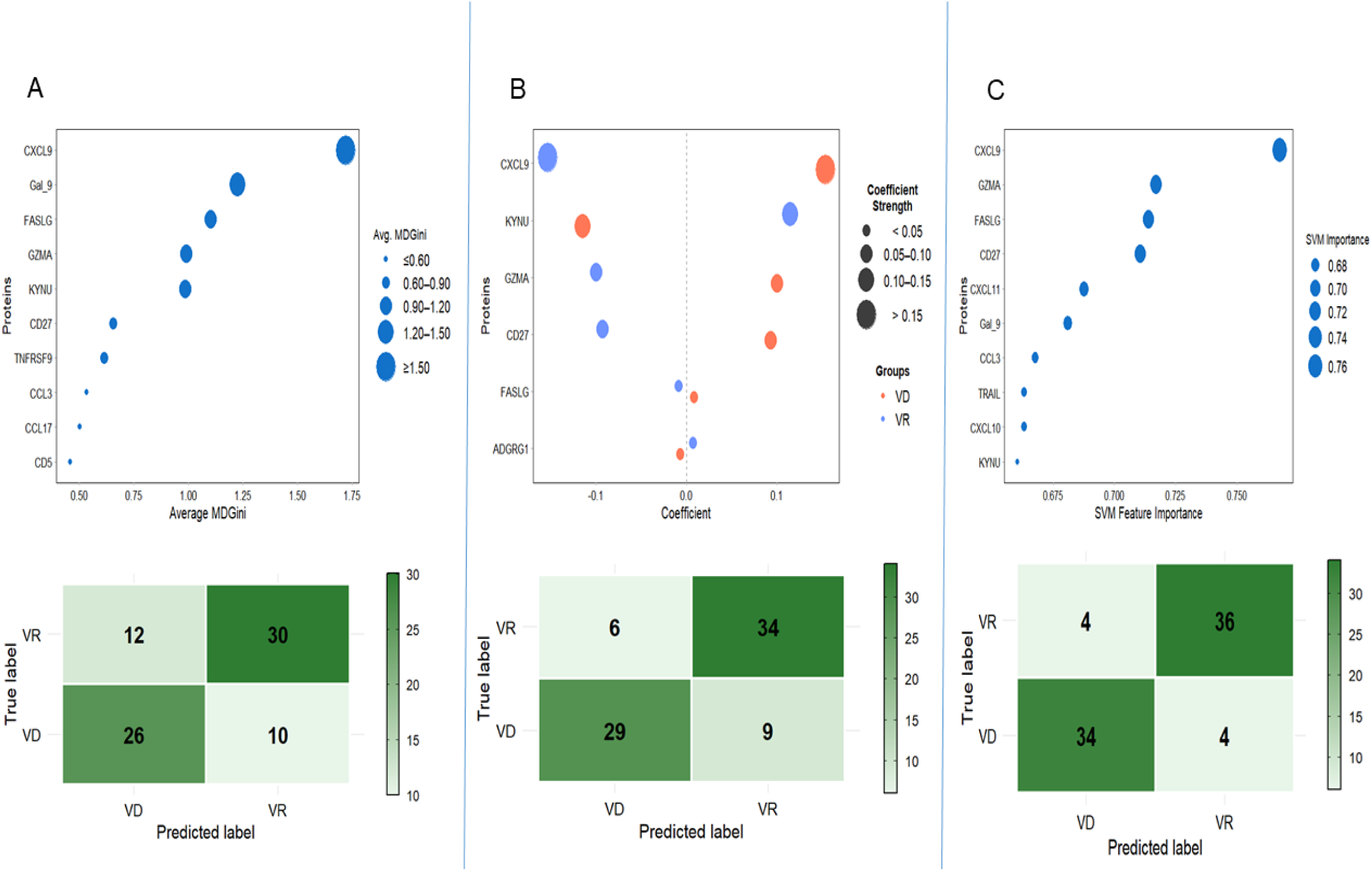
The most discriminative proteins selected by each model. Upper charts show top proteins selected by A: RF using Mean Gini Decreasing score, B: LASSO using L1-regularization, C: SVM using a RFE feature selection. The lower parts show the confusion matrices. The proteins are labeled according to their gene names. VD: visit at diagnosis, VR: visit at remission, RF: Random Forest, LASSO: Least Absolute Shrinkage and Selection Operator, SVM-RFE: Support Vector Machine Recursive Feature Elimination.

The LASSO model employs L1-regularization to perform feature selection by applying a shrinkage penalty to the model coefficients. By penalizing the absolute magnitude of the coefficients, the model effectively performs automatic feature selection, resulting in an interpretable model. At the optimal value of λ (λ1se), six proteins (CXCL9, FASLG, CD27, KYNU, ADGRG1, and GZMA) were identified **(Figure 2B)**. Two proteins, KYNU and ADGRG1, showed an increase in abundance in VR, while the other four proteins had a decreasing trend.

According to the results of the SVM-RFE model, the proteins CXCL9, GZMA, FASLG, CD27, CXCL11, Gal_9, CCL3, TRAIL, CXCL10, and KYNU were the most effective in discriminating the VD and VR conditions from each other **(Figure 2C)**.

### Evaluation of potential biomarkers identified by ML algorithms

All three models were consistent in identifying five common proteins **(Figure 3)**, while each model suggested unique proteins as well.

**Figure 3.**
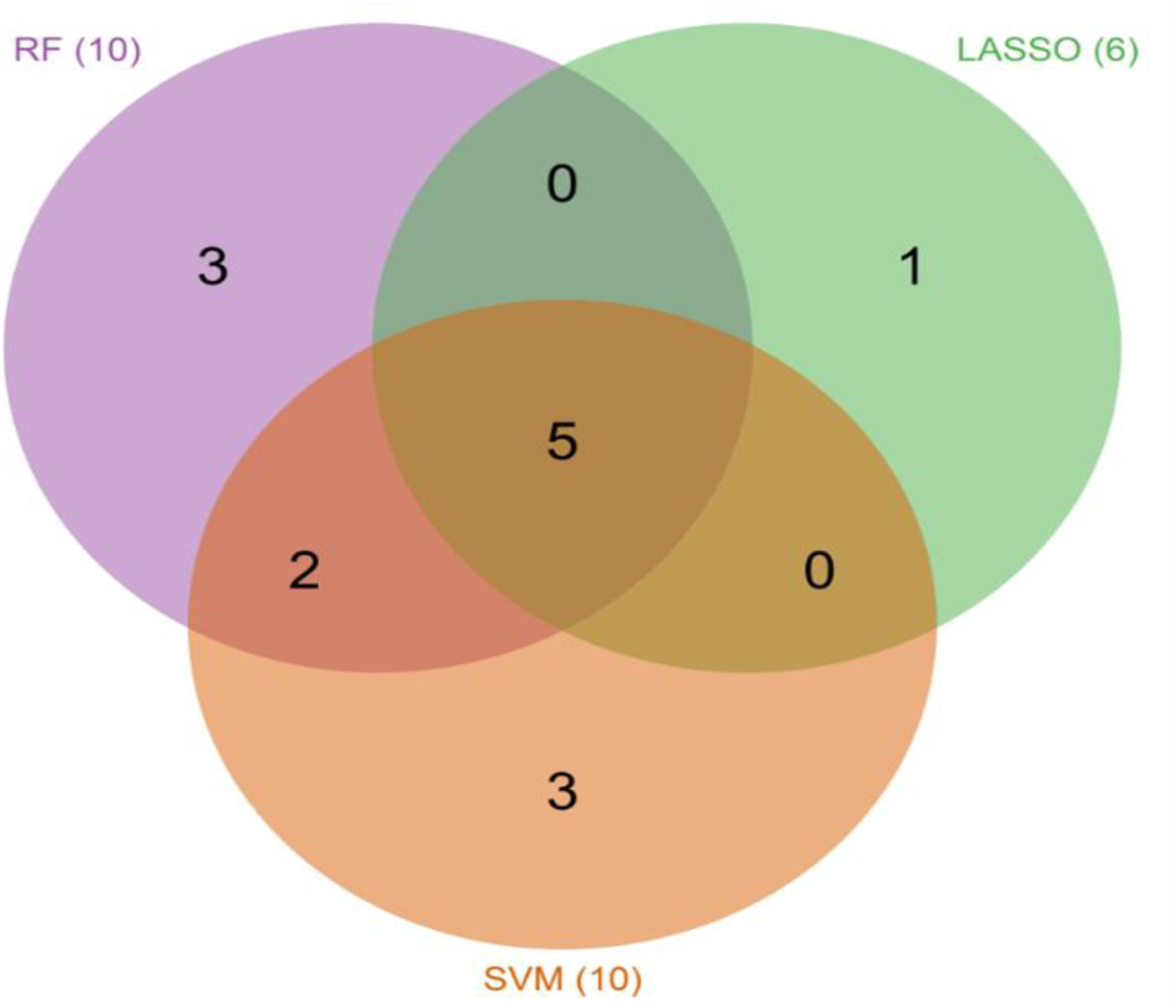
Comparative analysis of feature selection across RF, LASSO, and SVM-RFE models. The numbers in brackets refer to the number of proteins selected by each model. RF: Random Forest, LASSO: Least Absolute Shrinkage and Selection Operator, SVM-RFE: Support Vector Machine Recursive Feature Elimination.

According to the applied SVM model, the common five proteins performed well in the discrimination between the groups with an AUC of 0.775. We added each protein one by one to the model, and those that improved model performance were kept, and then, that combination of proteins was selected that provided the highest AUC **(Supplementary Table 3)**. According to our results, the combination of ten proteins: ADGRG1, CCL17, CD5, CD27, CXCL9, CXCL11, FASLG, GZMA, KYNU, and TNFRSF9 can serve as a potential biomarker panel in discriminating VD from VR with an AUC of 0.866. The heat map representation of the selected proteins highlights their discriminatory potential **(Figure 4, and Supplementary Figure 5)**, although only in the case of CD27 and CXCL9 was the difference previously shown to be statistically significant.

**Figure 4.**
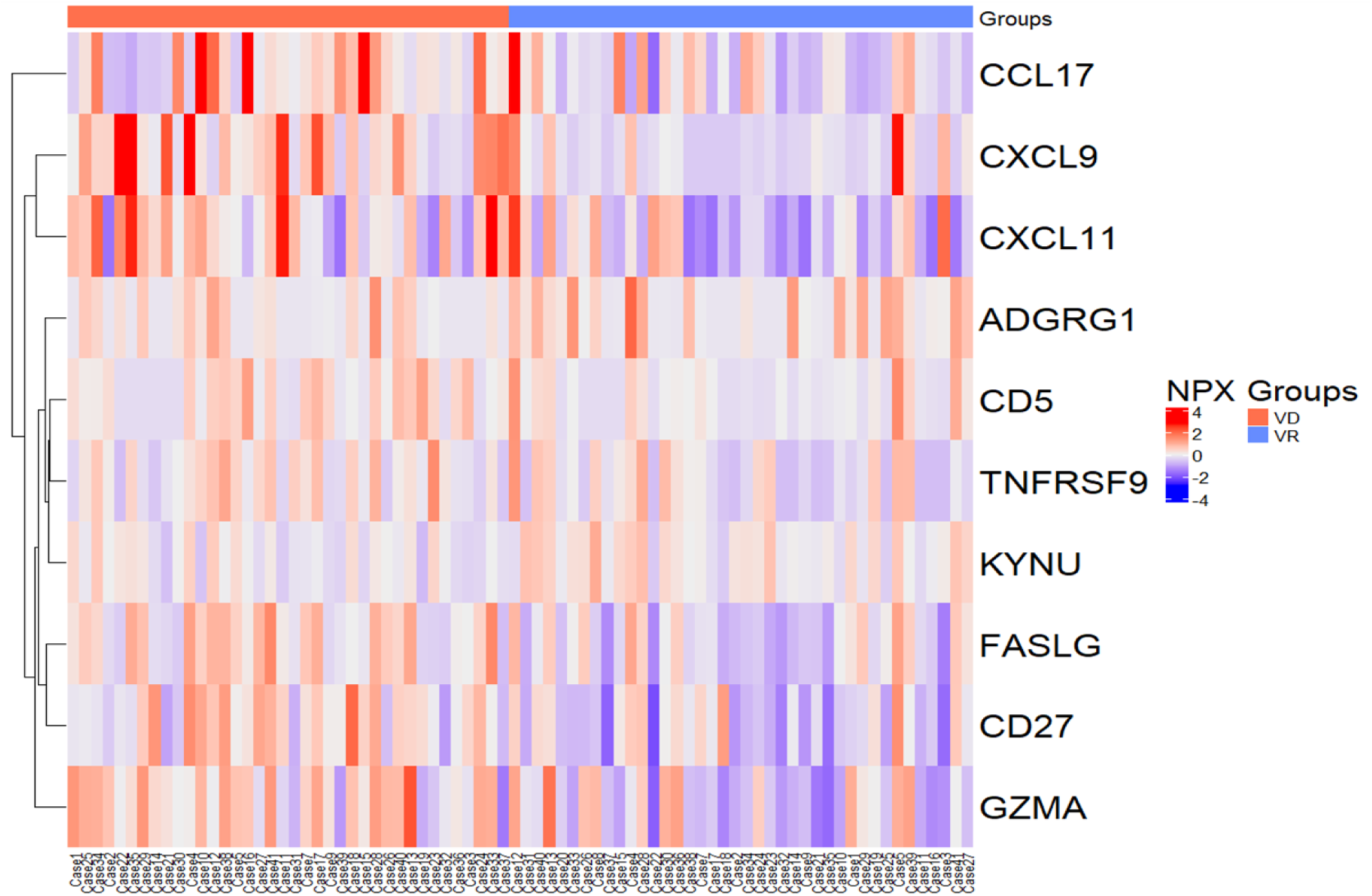
The heat map of the potential biomarkers in discriminating VD from VR. The columns represent the patients, and the rows represent the proteins. The colored bar above the graph indicates VD (orange) and VR (blue) groups, and red and violet in the NPX bar represent high and low NPX, respectively. VD: visit at diagnosis, VR: visit at remission, NPX: normalized protein expression

### Protein Expression Profiling Across Cancer Types

To prove the clinical relevance of the identified protein panel, we verified the expression of these proteins in the TPCPA dataset (n=999) [16]. Among the initial candidates, ADGRG1, CD5, KYNU, and GZMA demonstrated significant protein expression across various types of leukemia, like T-ALL or B-ALL. These markers are not uniformly expressed but show distinct patterns across different leukemia subtypes **(Supplementary Figure 6)**.

Some of the proteins including ADGRG1 and KYNU, were identified as predictive biomarkers in several cancer types, such as acute myeloid leukemia, melanoma, colon, breast, ovarian, esophageal squamous cell cancer, non-small cell lung, pancreatic, and glioblastoma/astrocytoma [21], [22], [23], [24].

### Survival Analysis of Upregulated and Downregulated Gene Signatures

Evaluation of the survival analysis within the Open Pediatric Cancer (OpenPedCan) Project (v15) revealed that lower expression of ADGRG1 and KYNU, as well as higher expression of CCL17, CD5, CD27, CXCL9, CXCL11, FASLG, GZMA, and TNFRSF9, showed no significant difference in survival between the altered and unaltered groups which were defined as having or not having alteration in queried genes in the selected profiles **(Supplementary Figure 7)**. However, when narrowing the dataset exclusively to ALL patients, a distinct trend emerged; although lower expression of ADGRG1 and KYNU was associated with a better overall survival rate, this change was not statistically significant. Conversely, higher expression of CCL17, CD5, CD27, CXCL9, CXCL11, FASLG, GZMA, and TNFRSF9 predicts a significantly poorer survival outcome **(Supplementary Figure 7)**.

### Functional analysis of potential biomarkers

The network of the selected proteins and their first shell interactors revealed a highly interconnected cluster and two smaller clusters **(Figure 5A)**.

**Figure 5.**
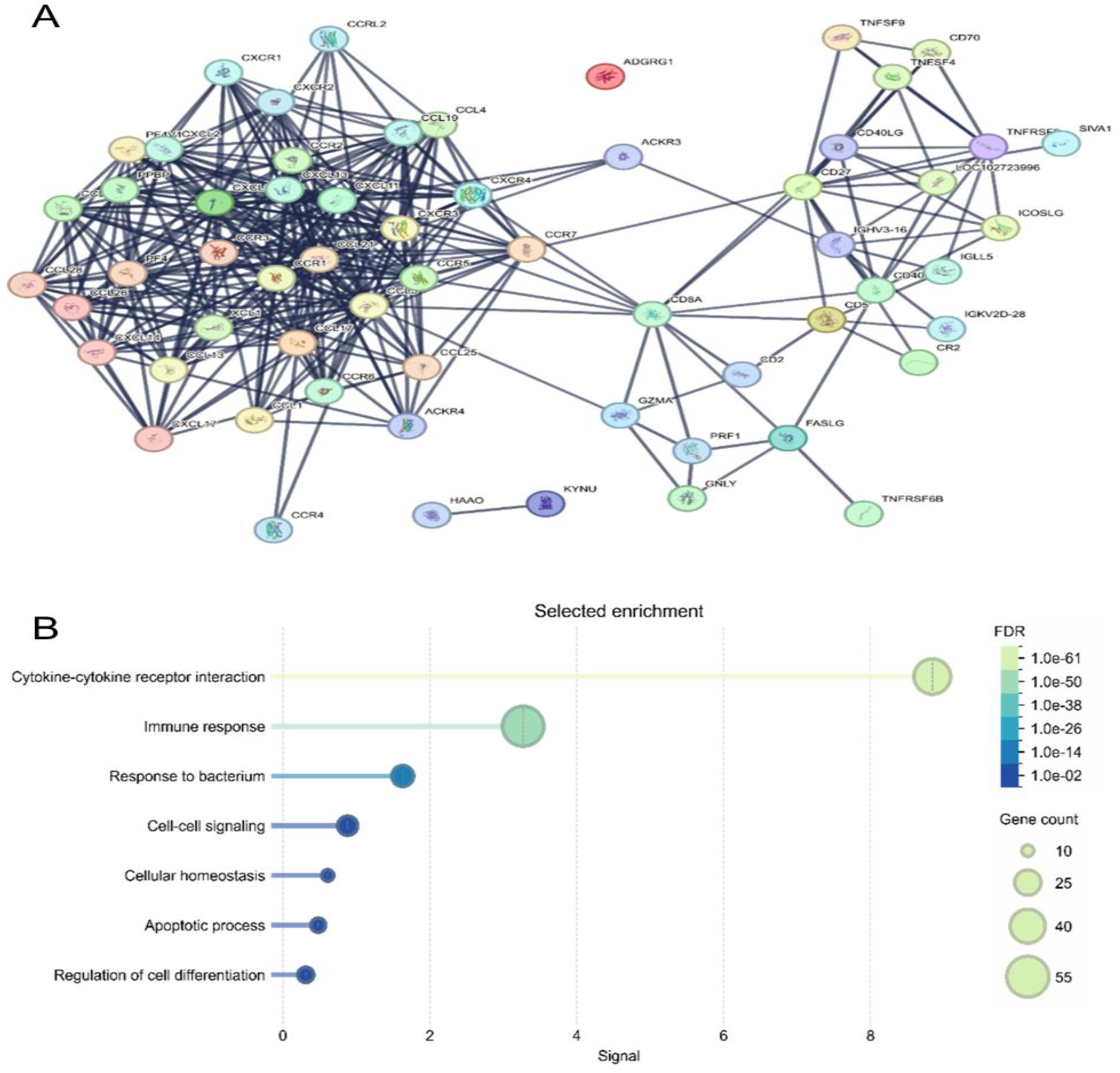
Protein-protein interactions network and functional analysis of the potential biomarkers. A: PPI network. Nodes represent proteins; edges represent protein-protein interactions, where the line thickness corresponds to the strength of data supporting the interaction. B: Functional enrichment analysis. The selected enriched pathways are shown; the colors indicate FDR p value, while the size of the circles represents the gene counts. FDR: false discovery rate.

The functional analysis indicated the involvement of immune function-related pathways, such as cytokine-cytokine receptor interaction, immune response, and response to bacteria. The cell-cell signaling, cellular homeostasis, apoptotic process, as well as regulation of cell differentiation indicate the involvement of complex mechanisms at the level of CSF in the changes related to remission **(Figure 5B)**.

### CSF Protein Profiles Discriminate CNS Involvement Levels Across VD and VR

To identify proteins capable of discriminating between patients with CNS involvement level 1 from other CNS involvement levels, we employed the same three different ML models. The models identified CCL4, CTSC, CXCL10, CXCL9, and MMP7 as discriminative features at the VD stage, while CAIX, CASP-8, HAGH, CXCL9, MMP7, MCP-2, and VWC2 were highlighted as significant discriminators at the VR stage.

At VD, CXCL9 and CXCL10 displayed elevated expression profiles in the CNS1 group, suggesting their potential role in early-stage disease regulation. Conversely, MMP7 exhibited heterogeneous expression, with notable upregulation in a subset of patients within the CNS2+3 group while at VR, MMP7 and CXCL9 demonstrated markedly higher expression in patients with CNS>1 involvement levels compared to CNS1. Conversely, CAIX showed a profile more characteristic of the CNS1 group, with relatively lower expression in the advanced CNS involvement stages **(Supplementary Figure 8, Supplementary table 4)**.

## Discussion

Our results highlight the importance of changes in ten key proteins during remission compared to the time of diagnosis. We showed that the level of CCL17, CXCL9, CXCL11, CD27, CD5, FASLG, GZMA, and TNFRSF9 decreased during remission, while ADGRG1 and KYNU showed an increasing trend.

These markers fall into three broad groups: immunoreceptors on lymphoid/T cells: CD27, CD5, ADGRG1, FAS/FASLG, chemokines: CXCL9, CCL17, CXCL11, and enzymes: KYNU, GZMA [25].

ADGR1 and KYNU were two proteins that showed an increased amount at the VR in our analysis. The ADGRG1 was previously identified as a potential biomarker in a broad range of cancers, including breast, melanoma, esophageal squamous cell cancer, non-small cell lung, ovarian, pancreatic, colon, glioblastoma/astrocytoma, and acute myeloid leukemia [15], [21], [22]. ADGRG1 acts as a marker for tumor-reactive T cells in acute myeloid leukemia; its targeting may help tumor-reactive TCR enrichment [26]. Transcriptomic data in Alzheimer’s disease show that ADGRG1 drives a protective microglial phenotype. It achieves this by activating the MYC transcription factor, which upregulates genes responsible for cellular homeostasis, phagocytosis, and lysosomal functions [27].

KYNU can be considered as a dual target, first as a diagnostic, prognostic, and predictive marker in solid tumors like gastric cancer, and second as a treatment target [23], [24]. In gastric cancer, its high expression causes an unfavorable immune microenvironment and low survival rate [23]. In addition, in breast cancer with metastasis, the activity of KYNU was higher in comparison to healthy individuals [28]. Different CNS disorders are linked to altered tryptophan metabolism, which primarily occurs through the kynurenine pathway. The resulting metabolites have diverse effects, including regulating neuronal excitability and triggering immune tolerance [29].

In contrast, a group of proteins, including chemokines like CCL17, CXCL9, CXCL11, and also CD27 and TNFRSF9, was decreased in VR. These proteins have been previously identified as potential biomarkers at the diagnosis and prognosis levels in T-cell and B-cell ALL and breast cancer [30], [31], [32], [33]. Increased expression of CXC chemokines, TNFRSF9, and CD27 was associated with poor prognosis and survival [32], [33], [34]. Our clinical verification using the OpenPedCan dataset strongly corroborates these findings specifically within the lymphoid lineage. While the 8-gene panel (CCL17, CD5, CD27, CXCL9, CXCL11, FASLG, GZMA, and TNFRSF9) demonstrated no prognostic impact in the heterogeneous pan-cancer population, their higher expression is associated with a significantly poorer survival outcome in ALL patients. This highlights that the observed decrease of these proteins during remission reflects a clinically favorable environment. Surprisingly CD27 showed a statistically significant decrease only in female patients, pointing toward a gender bias. According to the available data, the expression of CD27 associated with estrogen levels that showed a positive correlation with the frequency of CD27 negative naïve B cells, and a negative correlation with the frequency of CD27 positive memory B cells [35]. At the same time, soluble CD27 was found to be a promising CSF biomarker for tracking T-cell activation within the CNS [36]. The levels of CXC chemokines in patients with cancer, including breast, gastric, prostate, and colorectal cancer, can change dynamically depending on the tumor status, for example, whether the tumor is recurrent or resistant to chemotherapy. Accordingly, there is a potential to use these chemokines as diagnostic and prognostic biomarkers [32].

There is some evidence that CD5, FASLG, and GZMA proteins can also be introduced as potential biomarkers for some cancer types, including diffuse large B-cell lymphoma, breast cancer, small cell lung cancer, and ALL [37], [38], [39], [40]. Using bioinformatics on RNA-seq data from 1086 breast cancer patients from the TCGA database to assess the tumor microenvironment, CD5 was identified as a hub gene. High expression of CD5 is significantly associated with increased survival rates; its expression was associated with increased infiltration of cytotoxic CD8 positive T cells and M1 macrophages, and decreased presence of M2 macrophages [41]. Elevated CD5 levels indicate inflammation and cellular injury [42]. It is a circulating biomarker to cerebral palsy risk, establishing it as a potential target for both pharmaceutical development and lifestyle modifications to improve disease prophylaxis and care [43].

Chemokines and their receptors play a critical role in attracting peripheral white blood cells into the CNS, which drives the development of multiple sclerosis. Clinical, pathological, and brain imaging data indicate that the forms of this disease characterized by relapses—specifically the relapsing-remitting and secondary progressive variants—differ mechanistically from the primary progressive form [44]. For example, CXCL10 expression is prominent during CNS immunopathology, where it is believed to act as a primary regulator of T-cell-driven inflammation [45].

Similar findings were observed for FAS; patients with higher FAS expression exhibited a lower risk of progression from myelodysplastic syndrome to secondary acute myeloid leukemia [46]. While increased expression of GZMs serve as favorable prognostic indicators in urothelial carcinoma and cutaneous melanoma [47], they exhibit an opposite trend in renal cell carcinoma, where the higher level of GZMs is associated with worse overall survival [48].

In addition to proteins associated with remission status, our ML evaluation identified distinct proteomic signatures related to CNS involvement. CXCL9 and MMP7 were consistently selected as discriminative features at both VD and VR. Notably, CXCL9 emerged as a significant feature in both the remission kinetics analysis and the CNS involvement level discrimination models, underscoring its dual relevance in disease monitoring, however, this time it was elevated in patients with CNS2+3 disease.

Matrix metalloproteinase (MMPs) have been recognized as key drivers in the advancement and metastatic potential of various malignancies [49]. In the context of childhood ALL, the MMP7 has been identified as a genetic susceptibility factor. Notably, the A-181G polymorphism of the MMP7 promoter is associated with an elevated disease risk, particularly among male patients and those diagnosed at an age younger than 3.5 years [49]. MMP7 is very important in leukemia and acts beyond a simple tissue damaging enzyme. In fact, this enzyme is known in leukemia as a facilitator of invasion into distant tissues [50].

Although childhood ALL is characterized by complex alterations in protein-based signaling networks, uncovering these underlying dynamic changes at the protein level remains a significant challenge that cannot be achieved by genomic or transcriptome analysis alone [51]Detailed analysis of the proteins found in body fluids of cancer patients can give more information on the mechanisms driving malignant transformation. Considering our data in the One Health framework, we can provide with potential targets for future biomarker development for disease monitoring and management. Our findings highlight a coordinated reprogramming of immune and tumor-associated proteins within the CSF microenvironment during the transition from diagnosis to remission in ALL, emphasizing a gender bias in the case of some proteins. The identified signatures not only reinforce previously described prognostic and diagnostic roles but also underscore their potential utility in monitoring treatment response and guiding personalized therapeutic strategies shifting the focus toward the Precision One Health approaches.

## Limitations

The relatively small sample size and the uneven distribution of patients across the three groups in terms of CNS involvement are important limitations of our study. The lack of a healthy control group is another limitation, as obtaining CSF samples from children without a medical indication is not possible. This analysis was limited to proteins included in the pre-designed Olink panels, and we do not have information on other potential biomarkers. Besides these challenges, we also have to take into account the effect of chemotherapy that can severely affect the hepatic and renal functions of patients, manifesting as alterations in serum proteins. These systemic and metabolic fluctuations, alongside the proteins originating from damaged leukemic cells or therapy-induced immune responses, make it virtually impossible to isolate the true origin of proteomic changes when comparing post-treatment VD and VR samples. Finally, the scope of our external evaluation was restricted to extant datasets in public databases; consequently, the findings may reflect the specific characteristics of those study cohorts rather than the broader patient population. Future studies integrating larger longitudinal data and functional validation will be essential to clarify whether these proteins actively contribute to the molecular mechanisms of remission or serve primarily as indicators of systemic recovery, ultimately supporting their translation into clinical practice.

## Conclusion

In conclusion, the integration of Olink proteomics with ML algorithms identified distinct molecular signatures associated with remission and CNS involvement in ALL. These findings highlight the potential of high-dimensional proteomic profiling to improve disease monitoring and patient stratification. The observed sex-specific differences in the expression of several identified proteins underscore the need to account for gender as a critical biological variable when developing and validating biomarker panels. Future studies incorporating larger, independent cohorts and complementary validation approaches will be crucial for confirming these signatures and advancing their translation into clinical practice, ultimately supporting more precise, disease monitoring and fostering Precision One Health approaches.

## Abbreviations

ALL: Acute Lymphoblastic Leukemia
AUC: Area Under the Curve
CNS: Central Nervous System
CSF: Cerebrospinal Fluid
CV: Cross-Validation
FC: Fold Change
FDR: False Discovery Rate
LASSO: Least Absolute Shrinkage and Selection Operator
ML: Machine Learning
NPX: Normalized Protein Expression
OOB: Out-of-Bag
RF: Random Forest
SVM-RFE: Support Vector Machine Recursive Feature Elimination
VD: Visit of Diagnosis (initial visit)
VR: Visit of Remission (follow-up visit)

## Acknowledgements

This research was supported by the National Research, Development and Innovation Office(NKFIH) under grant number K143021. We are also grateful for the support provided by the Stipendium Hungaricum Scholarship. We thank Kamilla Sólyom for her excellent help in performing the Olink measurements, and Anna Emese Kiss for her invaluable assistance in CSF sample sorting.

## Author Contributions

Conceptualization: Éva Csősz, Csongor Kiss, Methodology: Mahshid Moballegh Nasery, Éva Csősz, Investigation and Data Collection: Réka Gergely, Nóra Borszékiné Kutszegi, Dániel János Erdélyi, Csongor Kiss, Istvan Szegedi, Mahshid Moballegh Nasery Resources: Csongor Kiss, Éva Csősz, Writing – Original Draft: Mahshid Moballegh Nasery, Éva Csősz, Writing – Review & Editing: Éva Csősz, Csongor Kiss, Dániel János Erdélyi. Supervision: Éva Csősz. All authors read and approved the final manuscript.

## Declaration of Generative AI and AI-assisted technologies in the writing process

During the preparation of this work, the author(s) used ChatGPT, Consensus, and Gemini in order to refine the structure of the manuscript, paraphrase some of the select sentences for clarity, and assist in developing code for statistical test selection based on data characteristics. After using these tools, the author(s) reviewed and edited the content as needed and take(s) full responsibility for the content of the published article.

