## Supplementary figure for "Unveiling Cerebrospinal Fluid Protein Biomarkers in Pediatric Acute Lymphoblastic Leukemia Using Proximity Extension Assay"

**Supplementary Figure 1**

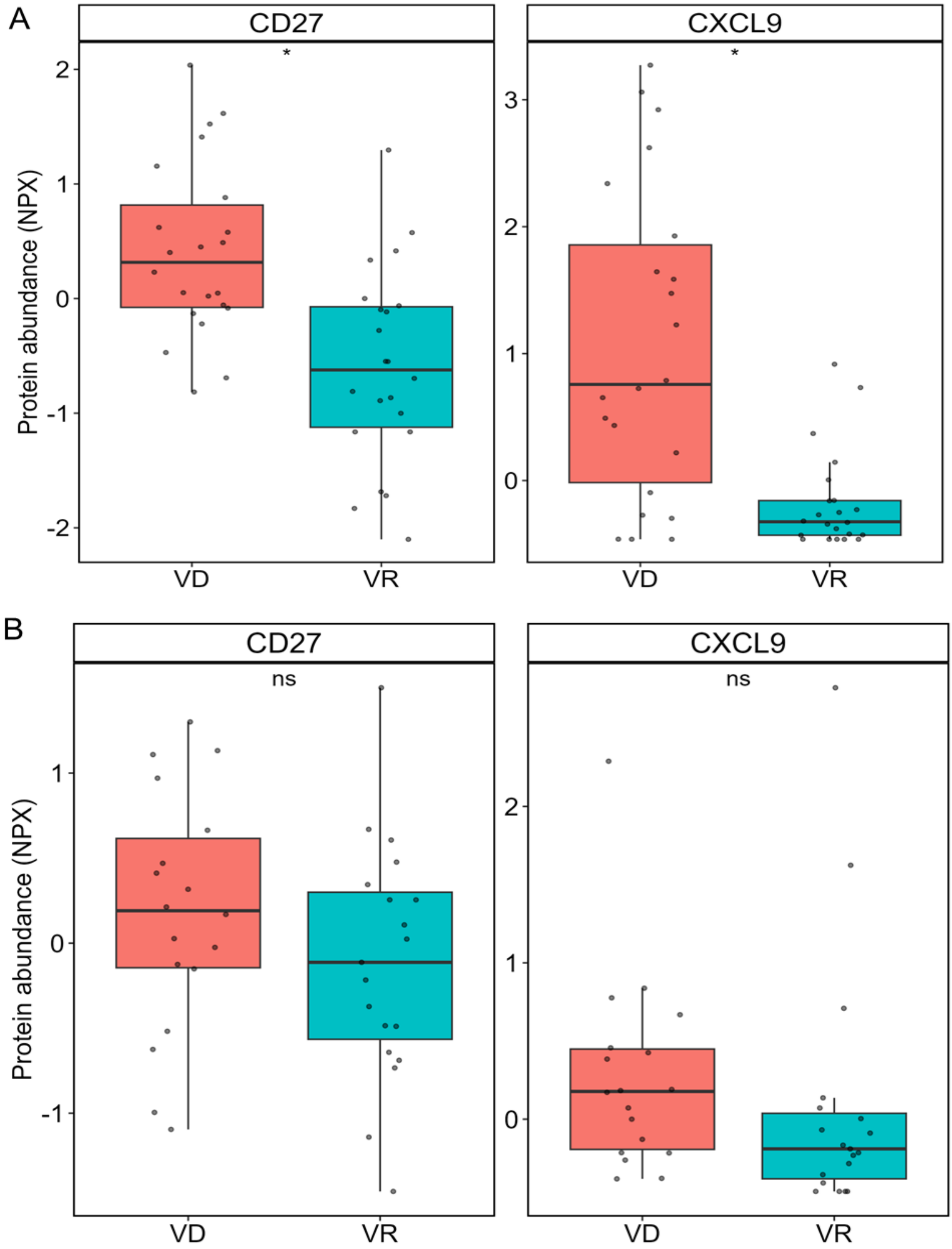


**Supplementary Figure 1.** Box plots of differentially abundant proteins in A: females and B: males. The boxes in red represent VD group, blue the VR group. VD: visit at diagnosis, VR: visit at remission

**Supplementary Figure 2**

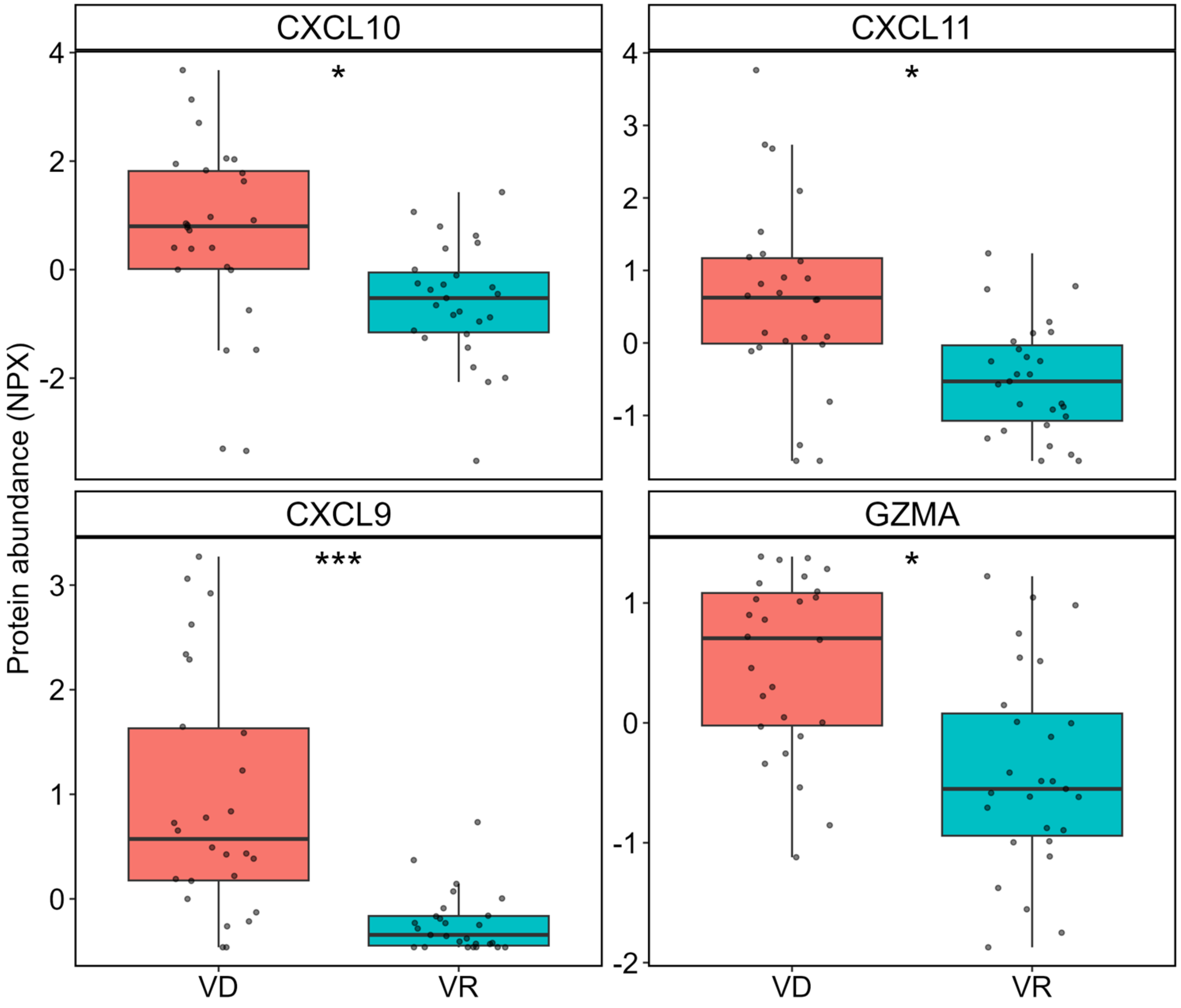


**Supplementary Figure 2.** Proteins showing statistically significant difference between the VD and VR groups in the CNS 1 group. The boxes in red represent VD group, blue the VR group.  VD: visit at diagnosis, VR: visit at remission

**Supplementary Figure 3**

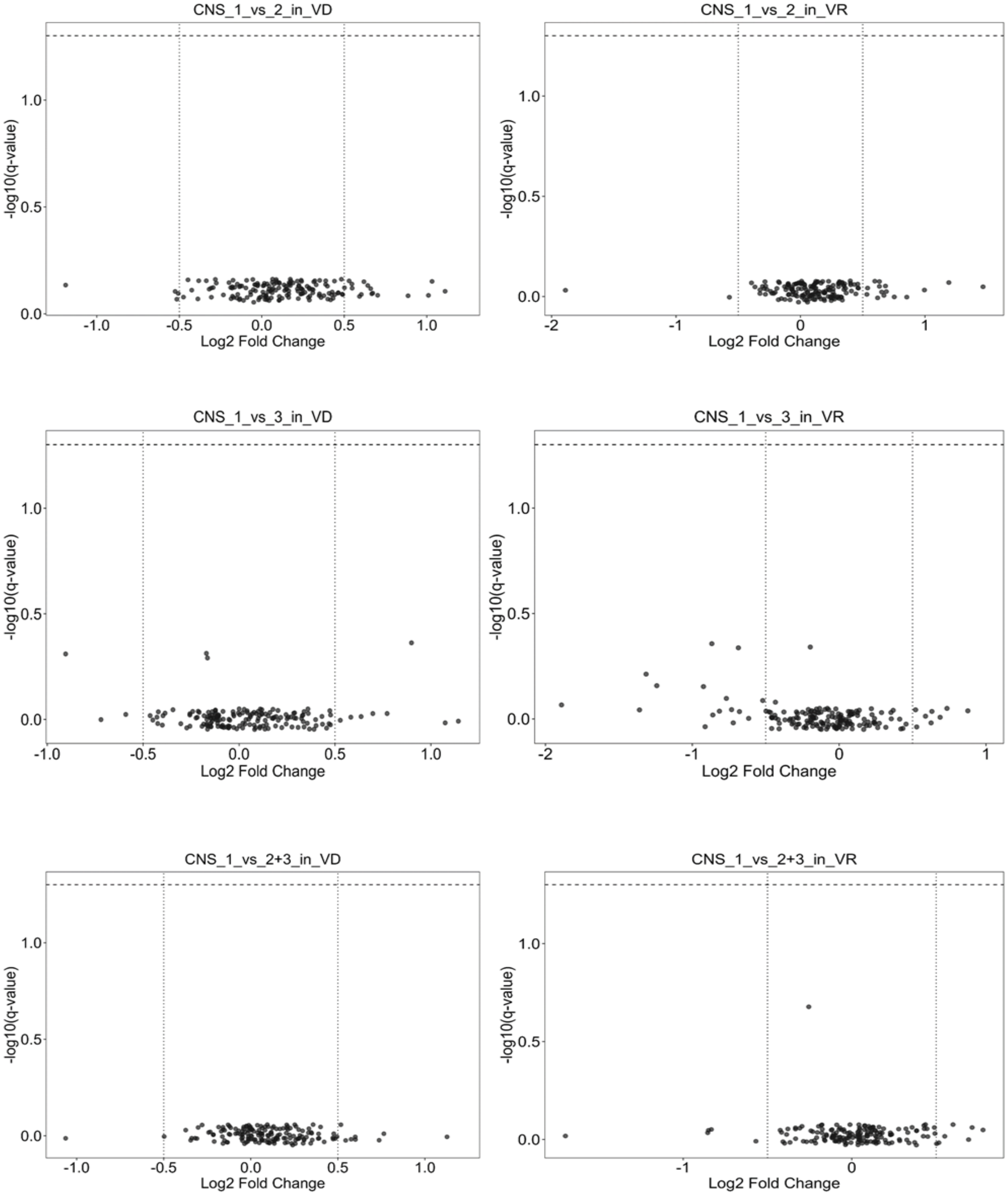


**Supplementary Figure 3.** Discrimination between CNS1 with other CNS variants across clinical time points. The figure displays comparisons of CNS1 versus CNS2, CNS3, and the combined CNS2+3 group, during both VD and VR. VD: visit at diagnosis, VR: visit at remission

**Supplementary Figure 4**

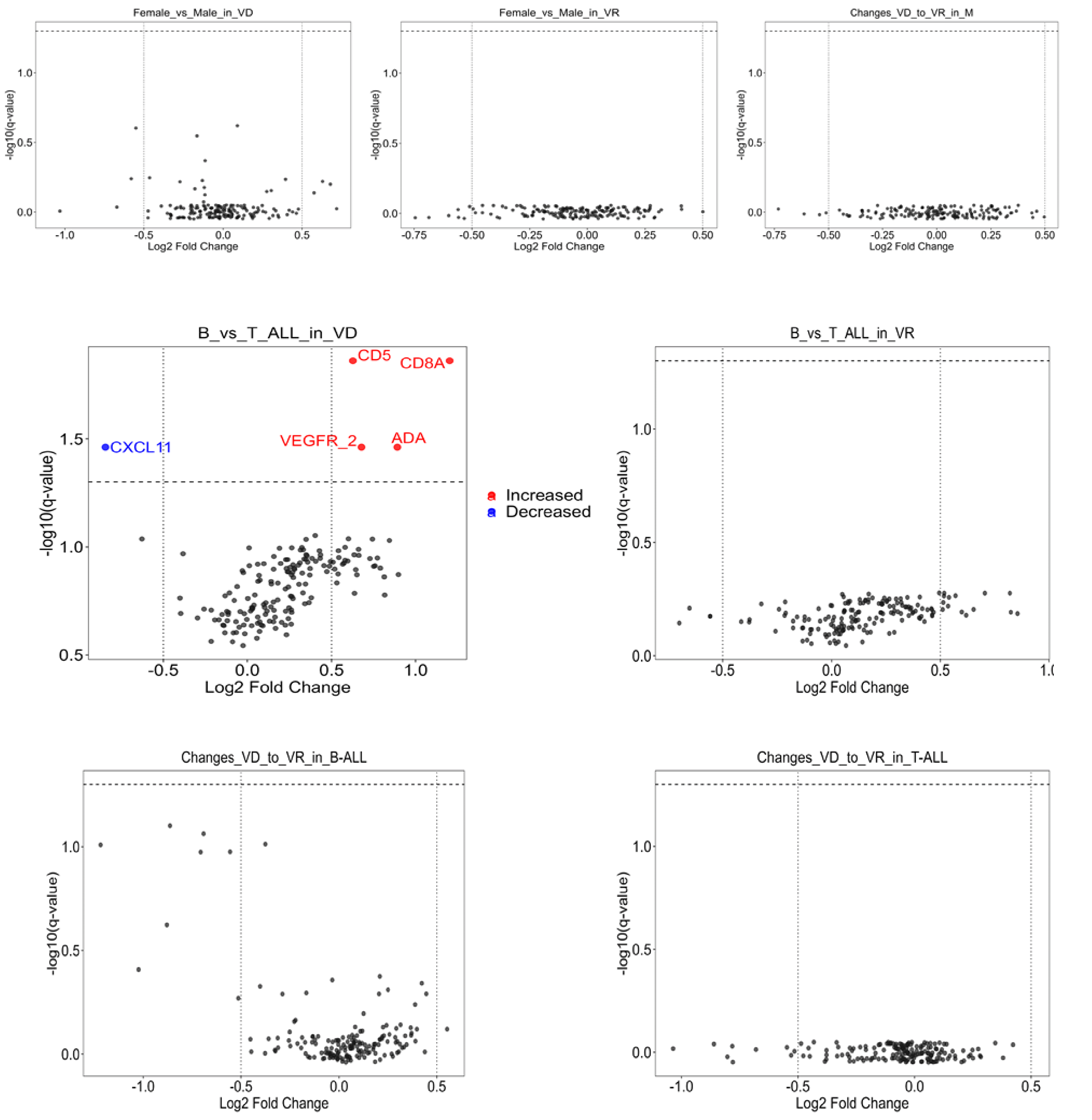


**Supplementary Figure 4.** Discrimination between of B and T-ALL and male and female across, the visit VD and the VR and within the visits. VD: visit at diagnosis, VR: visit at remission, ALL: Acute lymphoblastic Leukemia

**Supplementary figure 5**

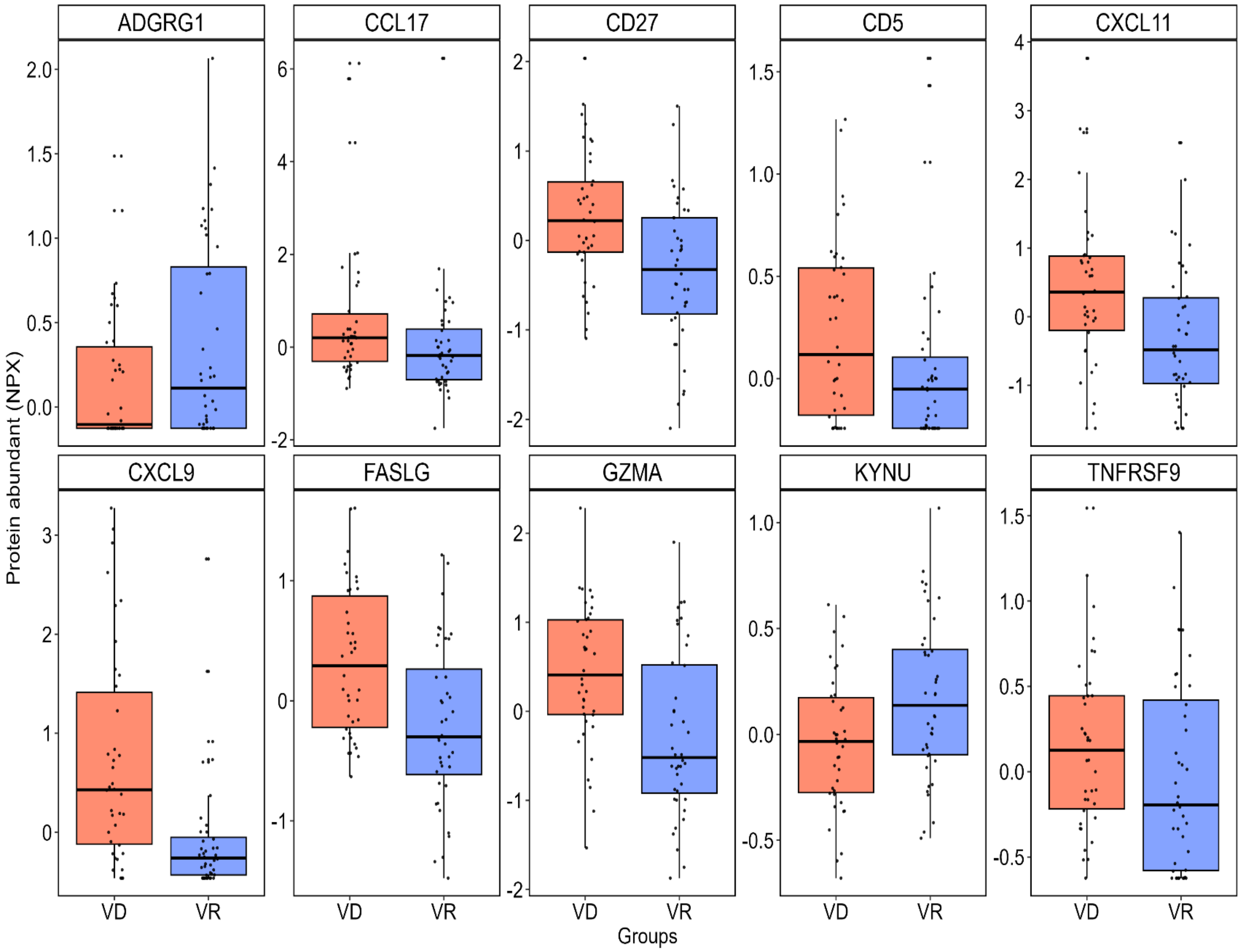


**Supplementary Figure 5.** Abundance changes of the selected potential biomarker proteins between the VD and VR groups. The boxes in red represent VD group, in blue the VR group. VD: visit at diagnosis, VR: visit at remission

**Supplementary figure 6**

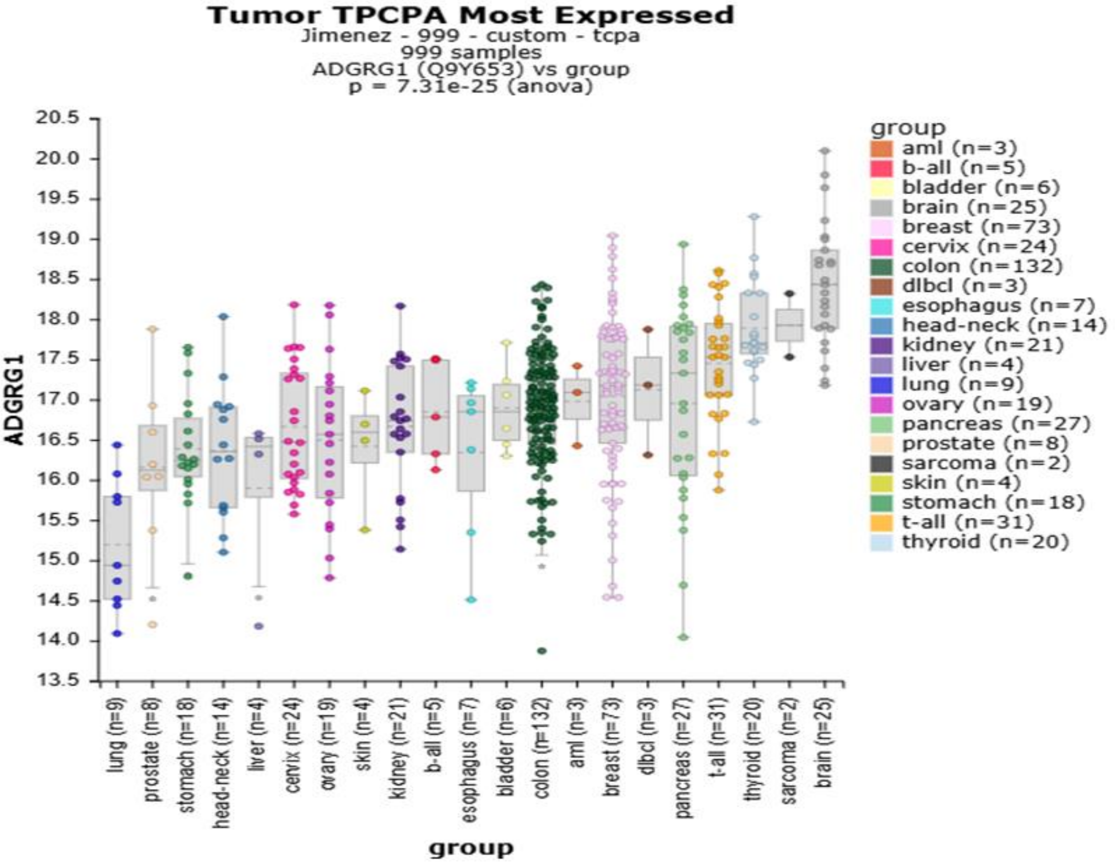


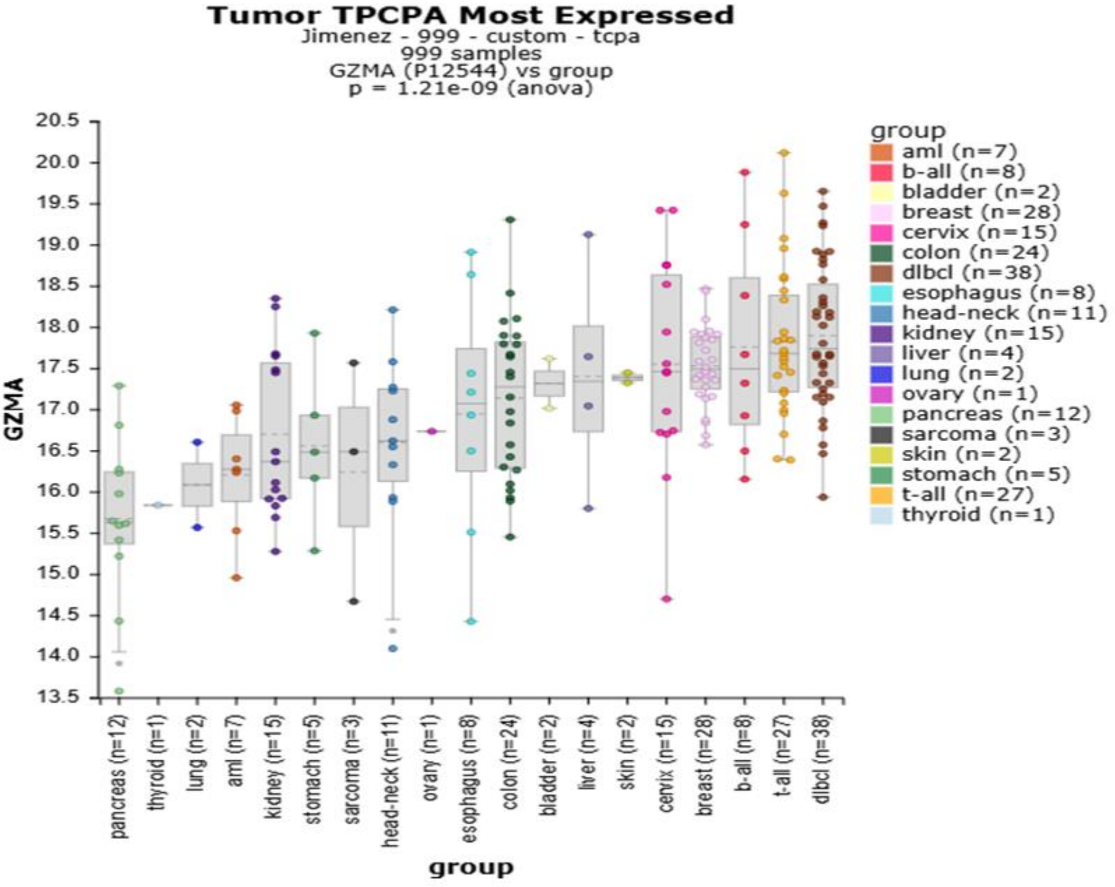

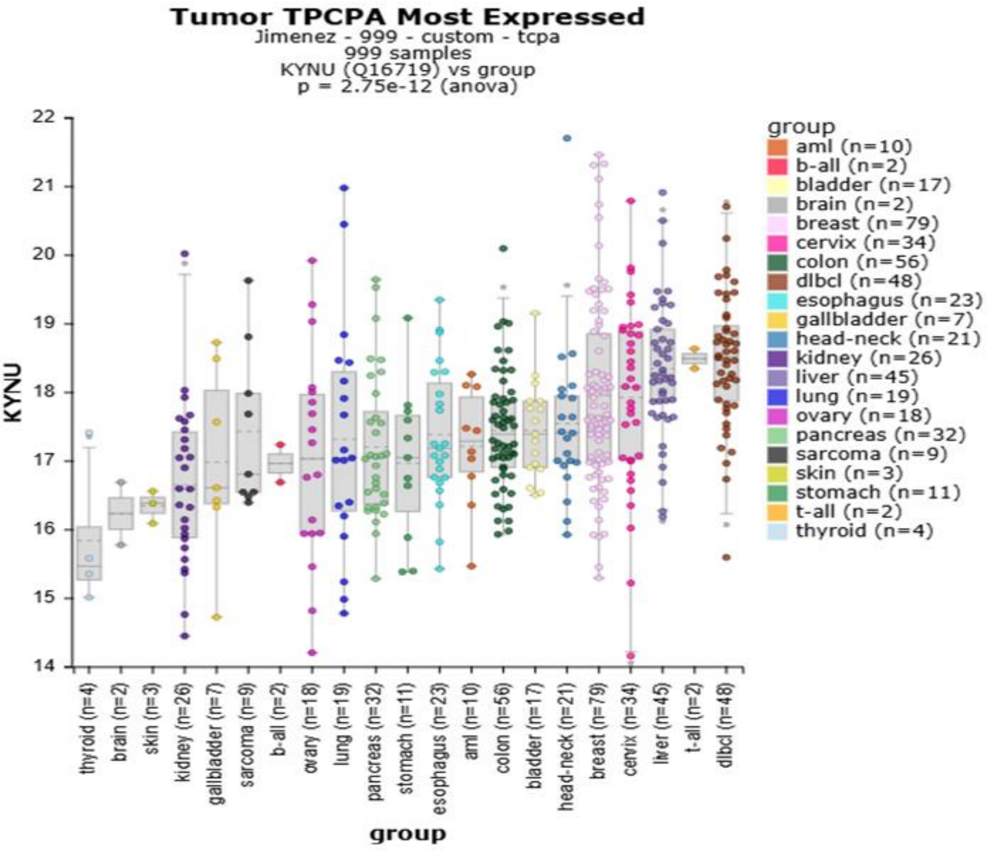
Tumor TPCPA Most Expressed
Jimenez - 999 - custom - tcpa
999 samples
KYNU (Q16719) vs group
p = 2.75e-12 (anova)
KYNU
group
thyroid (n=4)
brain (n=2)
skin (n=3)
kidney (n=26)
gallbladder (n=7)
sarcoma (n=9)
b-all (n=2)
ovary (n=18)
lung (n=19)
pancreas (n=32)
stomach (n=11)
esophagus (n=23)
aml (n=10)
colon (n=56)
bladder (n=17)
head-neck (n=21)
breast (n=79)
cervix (n=34)
liver (n=45)
t-all (n=2)
dlbcl (n=48)

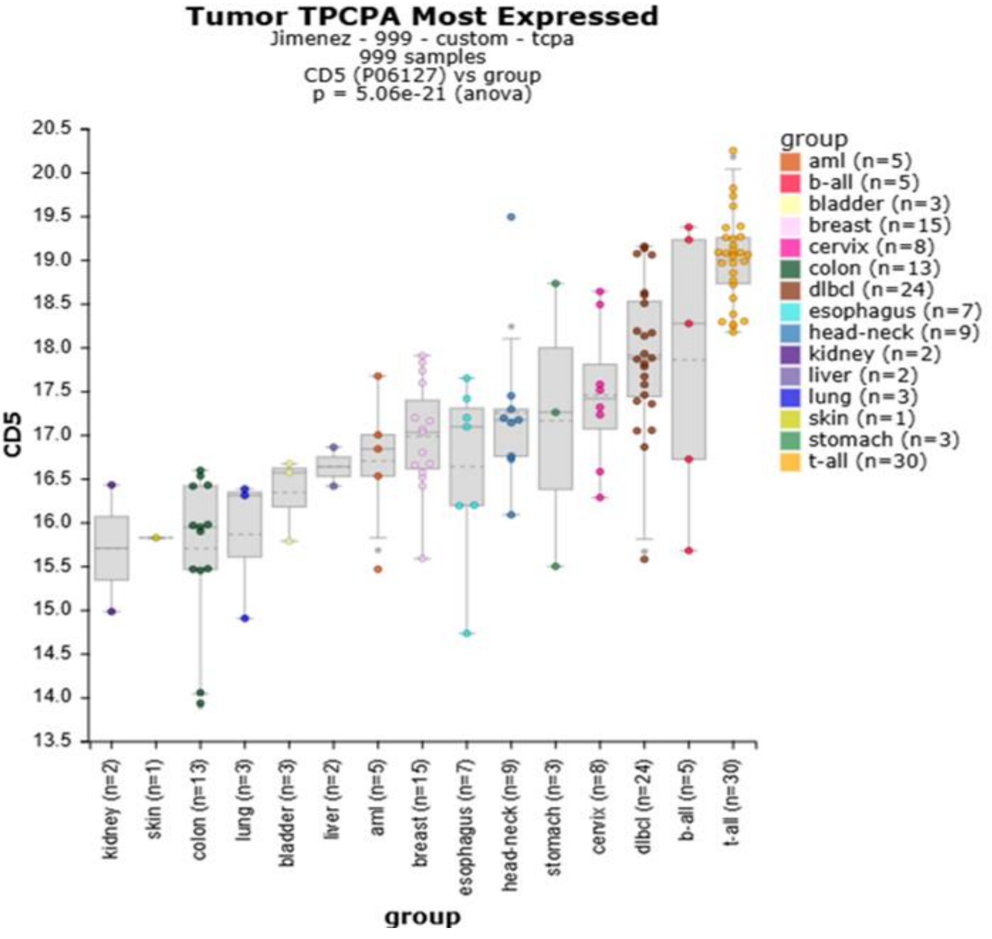
Tumor TPCPA Most Expressed
Jimenez - 999 - custom - tcpa
999 samples
CD5 (P06127) vs group
p = 5.06e-21 (anova)
CD5
group
kidney (n=2)
skin (n=1)
colon (n=13)
lung (n=3)
bladder (n=3)
liver (n=2)
aml (n=5)
breast (n=15)
esophagus (n=7)
head-neck (n=9)
stomach (n=3)
cervix (n=8)
dlbcl (n=24)
b-all (n=5)
t-all (n=30)

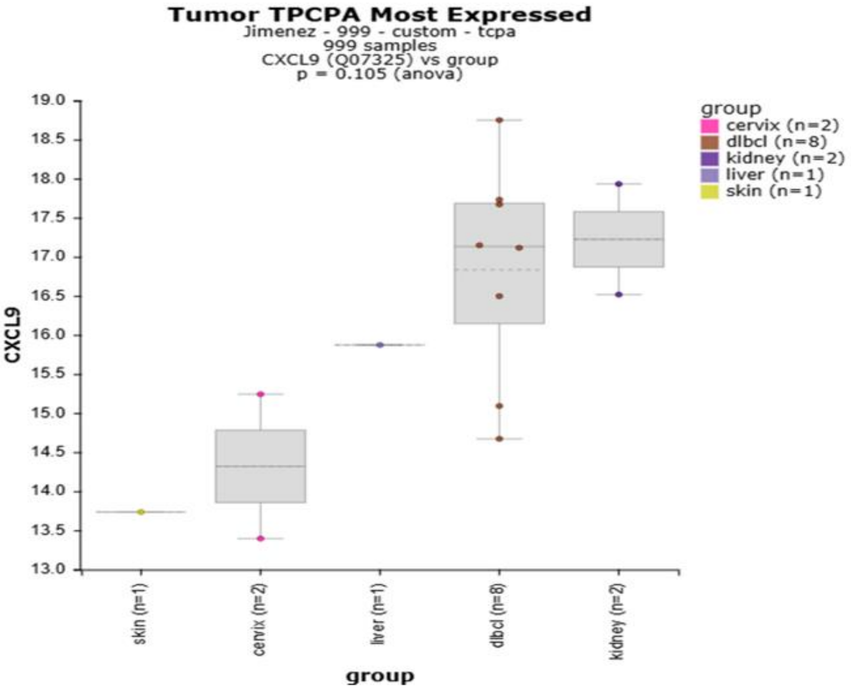


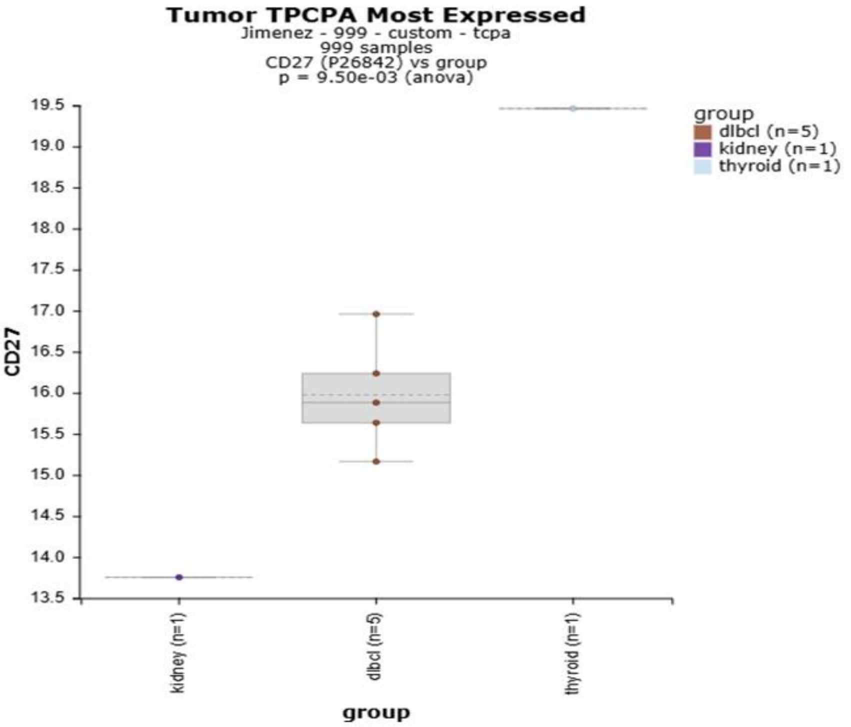


**Supplementary Figure 6.** Expression profiles of selected proteins in tumor types (Pan-Cancer).These graphs show the expression levels of CXCL9, CD27, CD5, GZMA, ADGRG1 and KYNU proteins in 31 different cancer types. The vertical axis shows the relative protein expression and the horizontal axis shows the different cancer groups. (The p-value indicated above each plot indicates the statistical significance of the difference between groups) (http://r2platform.com/TPCPA)

**Supplementary Figure 7**

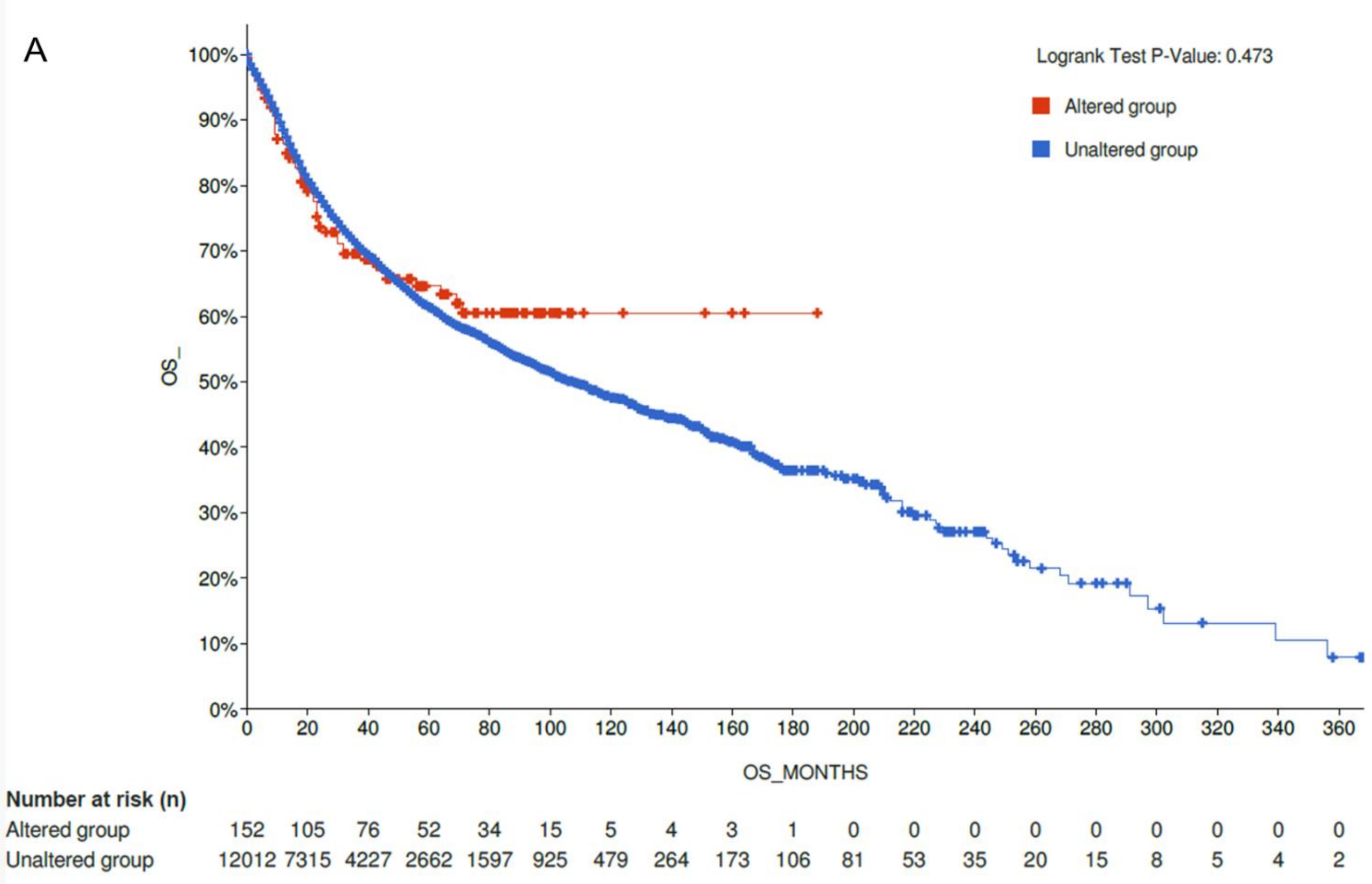

B

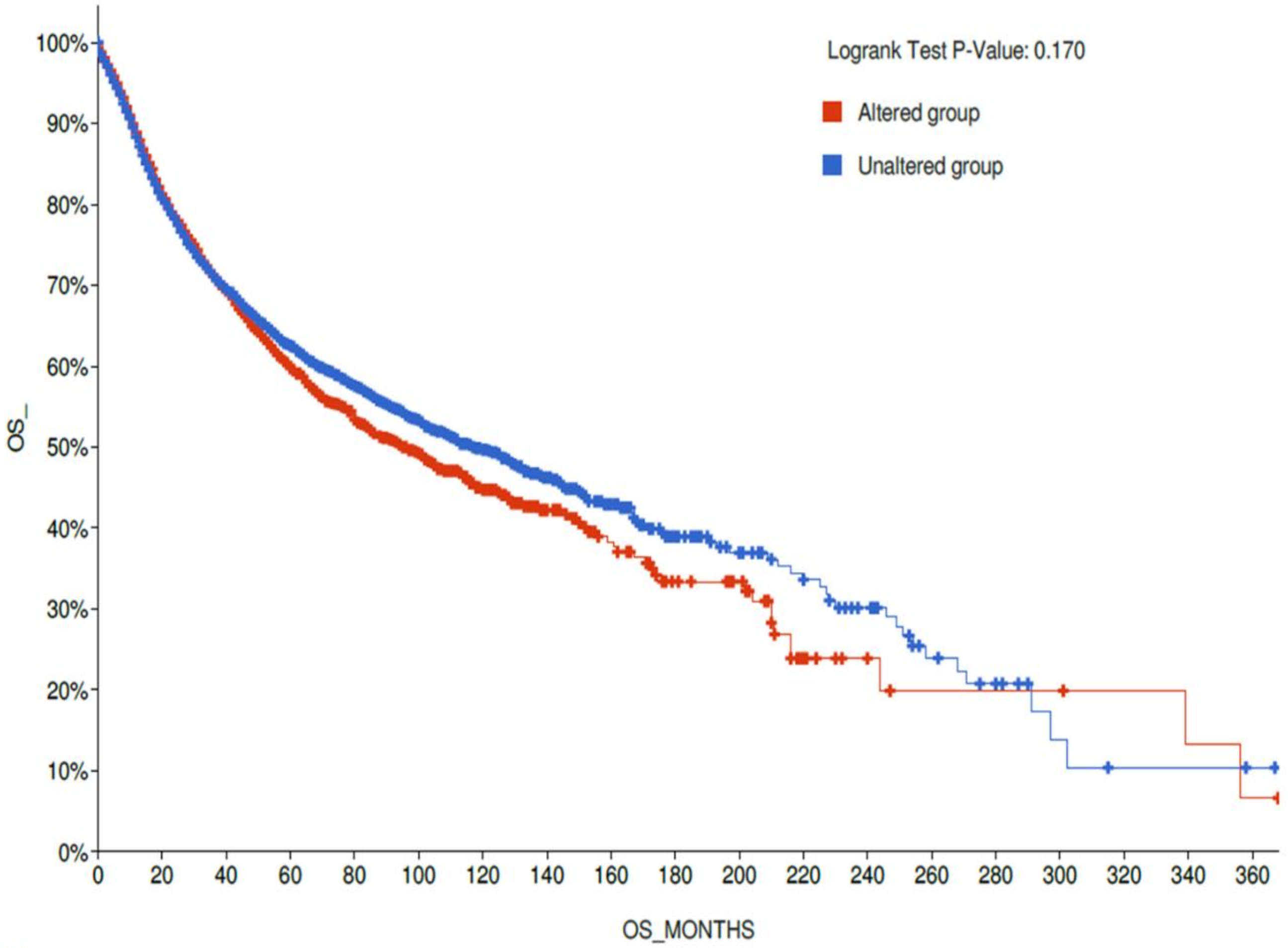


**Number at risk (n)**

| | | | | | | | | | | | | | | | | | | | |
|---|---|---|---|---|---|---|---|---|---|---|---|---|---|---|---|---|---|---|---|
| Altered group | 4076 | 2418 | 1267 | 741 | 428 | 265 | 156 | 90 | 62 | 35 | 30 | 13 | 7 | 4 | 4 | 4 | 3 | 2 | 1 |
| Unaltered group | 7823 | 4837 | 2941 | 1918 | 1177 | 670 | 322 | 179 | 116 | 74 | 52 | 41 | 29 | 16 | 11 | 4 | 2 | 2 | 1 |

C

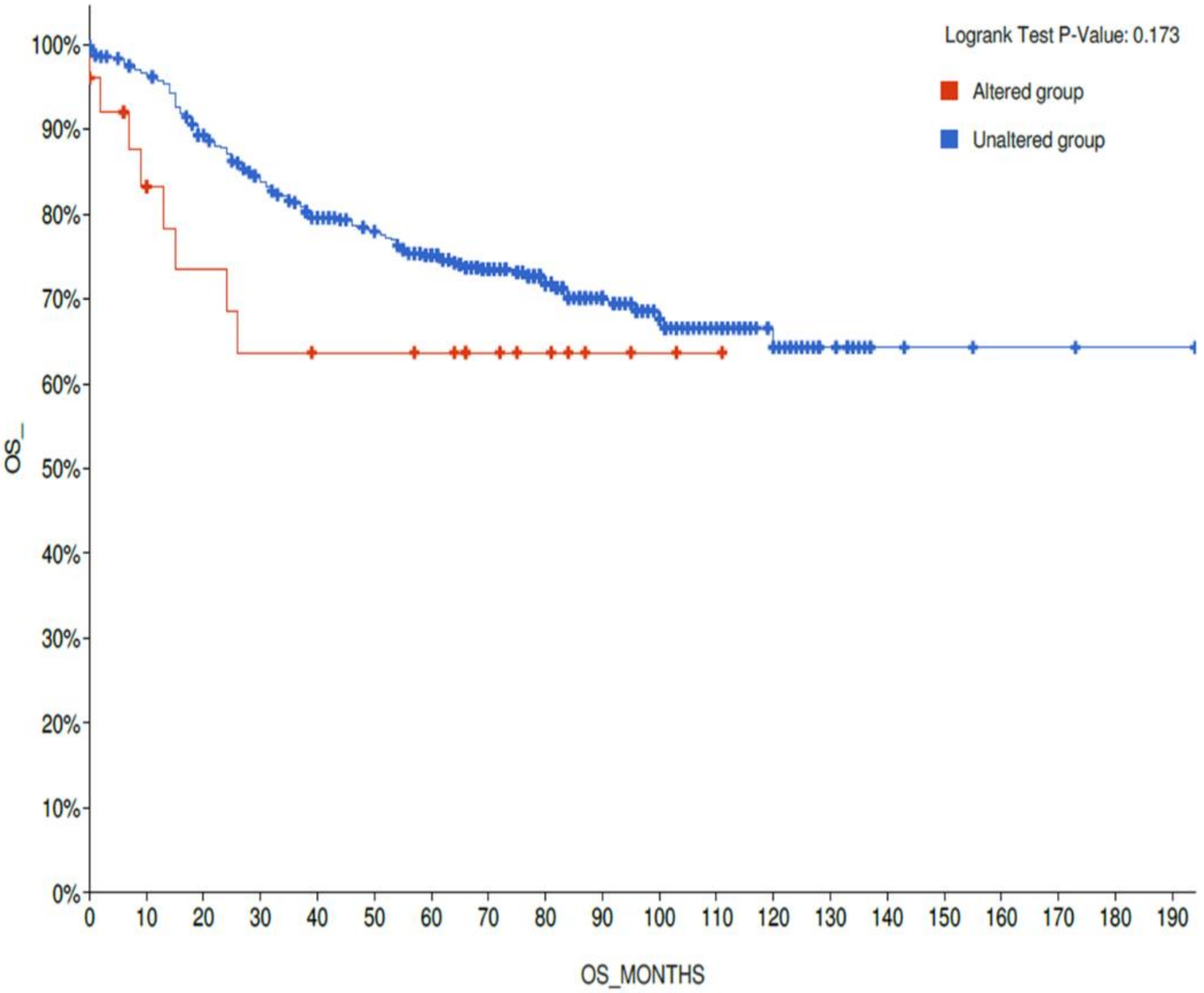


**Number at risk (n)**

| | | | | | | | | | | | | | | | | | | | | |
|---|---|---|---|---|---|---|---|---|---|---|---|---|---|---|---|---|---|---|---|---|
| Altered group | 25 | 19 | 15 | 13 | 12 | 12 | 11 | 8 | 6 | 3 | 2 | 1 | 0 | 0 | 0 | 0 | 0 | 0 | 0 | 0 |
| Unaltered group | 475 | 451 | 411 | 381 | 350 | 333 | 306 | 237 | 158 | 106 | 70 | 45 | 29 | 14 | 4 | 3 | 2 | 2 | 1 | 1 |

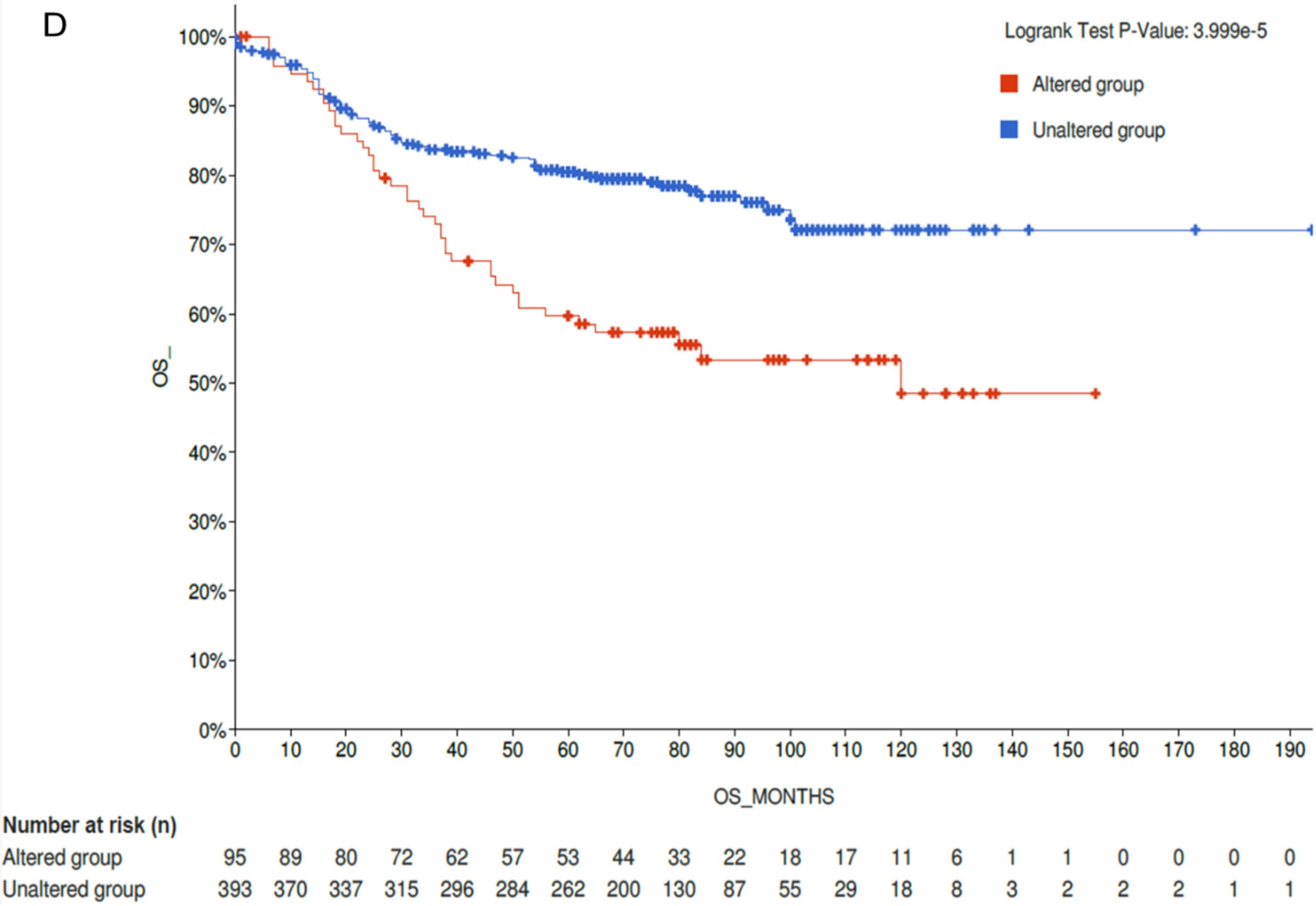


**Supplementary Figure 7.** Overall survival analysis, A: Lower expression of ADGRG1 and KYNU, and B: Higher expression of CCL17, CD5, CD27, CXCL9, CXCL11, FASLG, GZMA, and TNFRSF9 in Open Pediatric Cancer (OpenPedCan), C: Lower expression of ADGRG1 and KYNU, and D: Higher expression of CCL17, CD5, CD27, CXCL9, CXCL11, FASLG, GZMA, and TNFRSF9, narrowing the dataset exclusively to ALL patients from the same data set. ALL: Acute Lymphoblastic Leukemia, Altered group: sample with at least one alteration in queried genes in the selected profiles. Unaltered group: sample without any alteration in queried genes in the selected profiles. (https://pedcbioportal.org)

**Supplementary Figure 8**

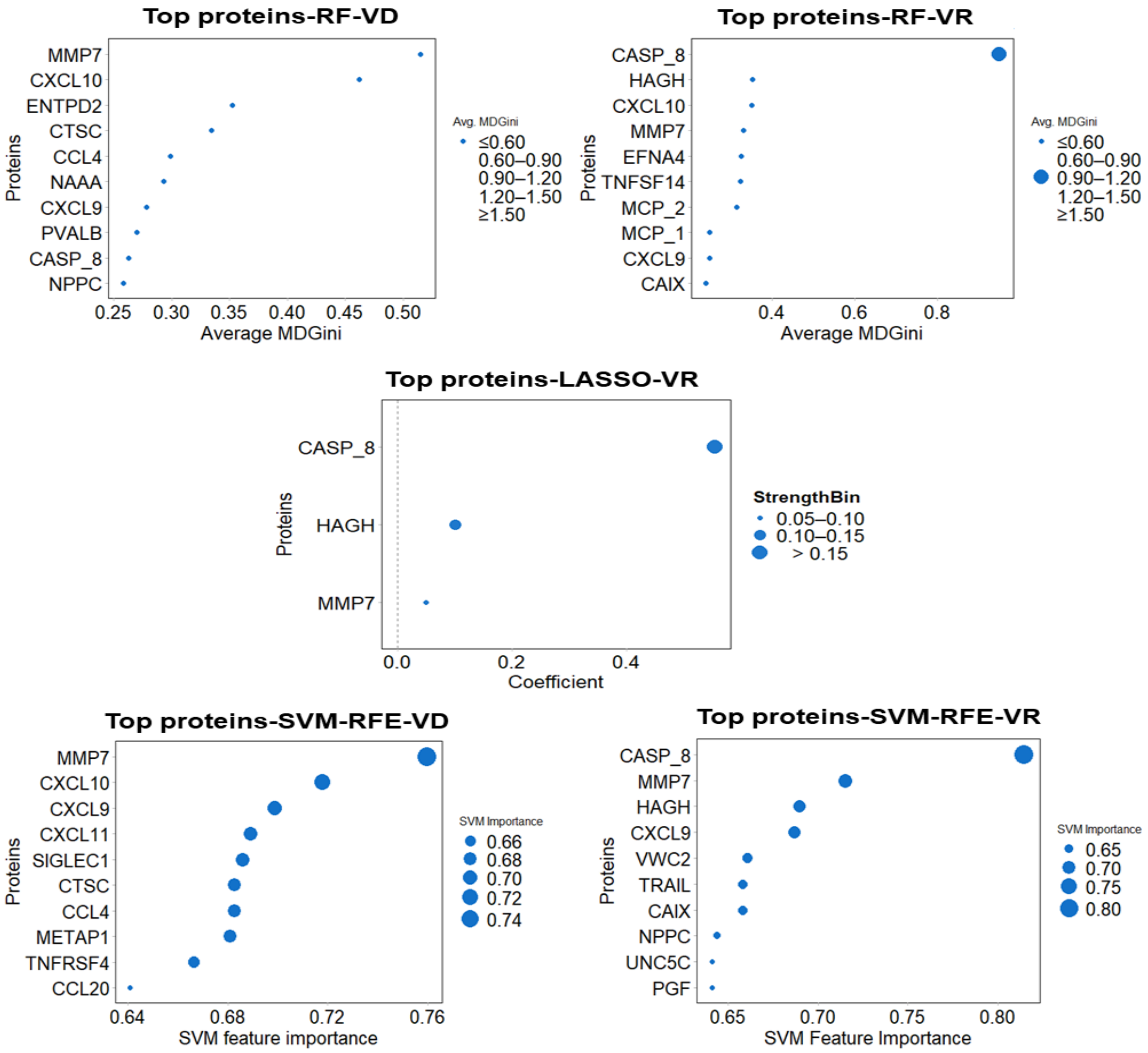

**Comarison among Three models (Rf/SVM/LASSO)_VD**

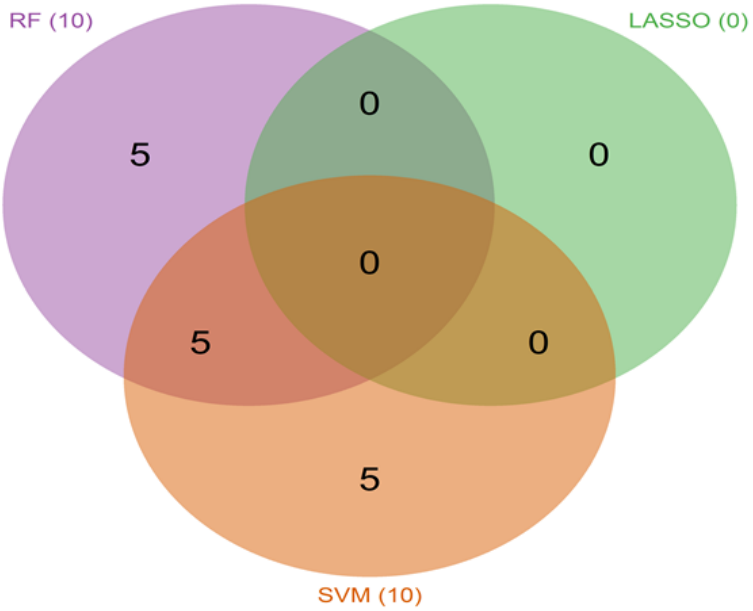


**Comarison among Three models (Rf/SVM/LASSO)_VR**

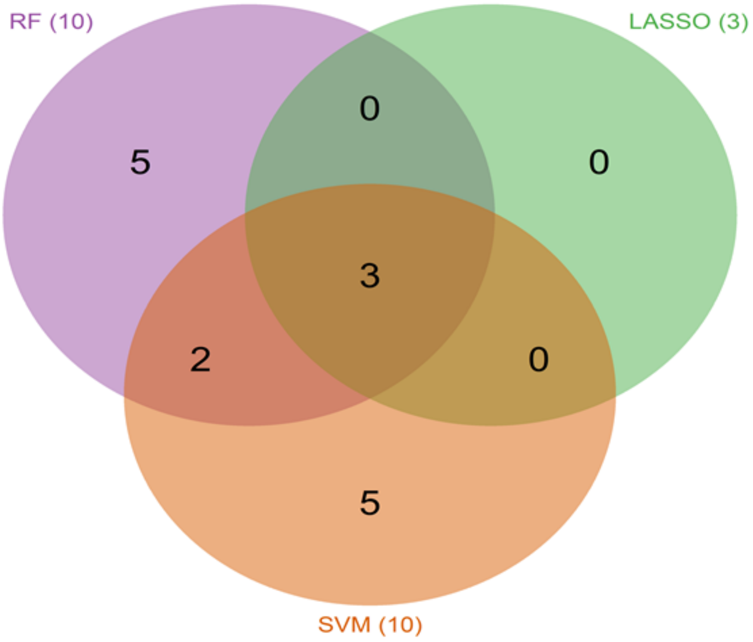

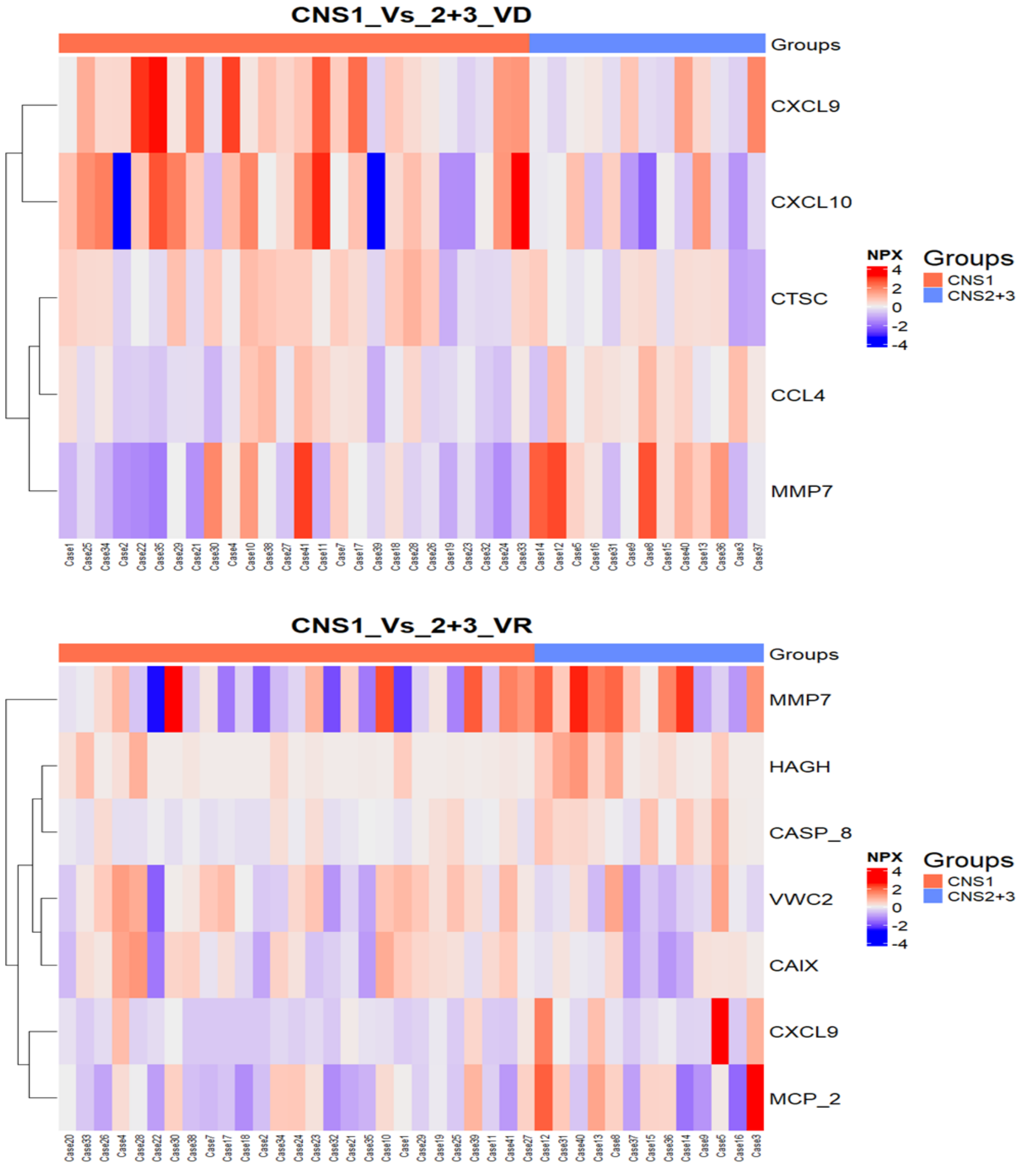


**Supplementary Figure 8**. Graphs shows the most important proteins selected by each model selected by RF using Mean Gini Decreasing score, LASSO using L1-regularization, and SVM using a RFE feature selection. Vendiagrams demonstrate comparative analysis of feature selection across RF, LASSO, and SVM-RFE models. The numbers in brackets refer to the number of proteins selected by each model. The heat map of the potential biomarkers in discriminating CNS1 from CNS2+3. The columns represent the patients, and the rows represent the proteins. The colored bar above the graph indicates VD (orange) and VR (blue) groups, and red and violet in the NPX bar represent high and low normalized protein expression, respectively. VD: visit at diagnosis, VR: visit at remission, RF: Random Forest, LASSO: Least Absolute Shrinkage and Selection Operator, SVM-RFE: Support Vector Machine Recursive Feature Elimination. NPX: normalized protein expression
